# *In vivo* BMAL1 occupancy mapping using MACS-Calling Cards reveals disease-associated retargeting in *Cln3^Δex7/8^* astrocytes

**DOI:** 10.64898/2026.04.30.721783

**Authors:** India H. Reiss, Jonathan D. Cooper, Erik S. Musiek, Robi David Mitra

## Abstract

Astrocytic homeostatic programs, many of which are regulated by the circadian clock, are disrupted early in neurodegenerative disease. The core clock transcription factor (TF) BMAL1 is required for normal astrocyte function, but its role during disease remains unclear. This is partly because methods for identifying cell type-specific TF binding sites are limited. Here, we developed MACS-Calling Cards (MACS-CC), a strategy for mapping astrocyte-specific TF occupancy *in vivo*. We used MACS-CC to define BMAL1 binding in the *Cln3^Δex7/8^* mouse model of CLN3 disease, a fatal neurodegenerative disorder marked by early astrocyte dysfunction and circadian disruption. BMAL1 binding was extensively redistributed in *Cln3^Δex7/8^* astrocytes: wild-type-specific binding sites enriched near glial differentiation genes, whereas *Cln3^Δex7/8^*-specific sites lacked functional enrichment. Consistent with these changes, *Cln3^Δex7/8^* astrocytes decreased expression of mature astrocyte markers. To define mechanisms underlying BMAL1 retargeting, we tested whether altered chromatin accessibility explained the changes in BMAL1 binding. Although chromatin accessibility was broadly remodeled, differential accessibility did not predict BMAL1 redistribution. Instead, motif analysis suggested that loss of cooperative TF interactions drives BMAL1 retargeting. These findings demonstrate that MACS-CC enables astrocyte-specific TF occupancy mapping and reveals mechanisms behind early rewiring of circadian regulatory programs within a model of a neurodegenerative disease.

**GRAPHICAL ABSTRACT:** 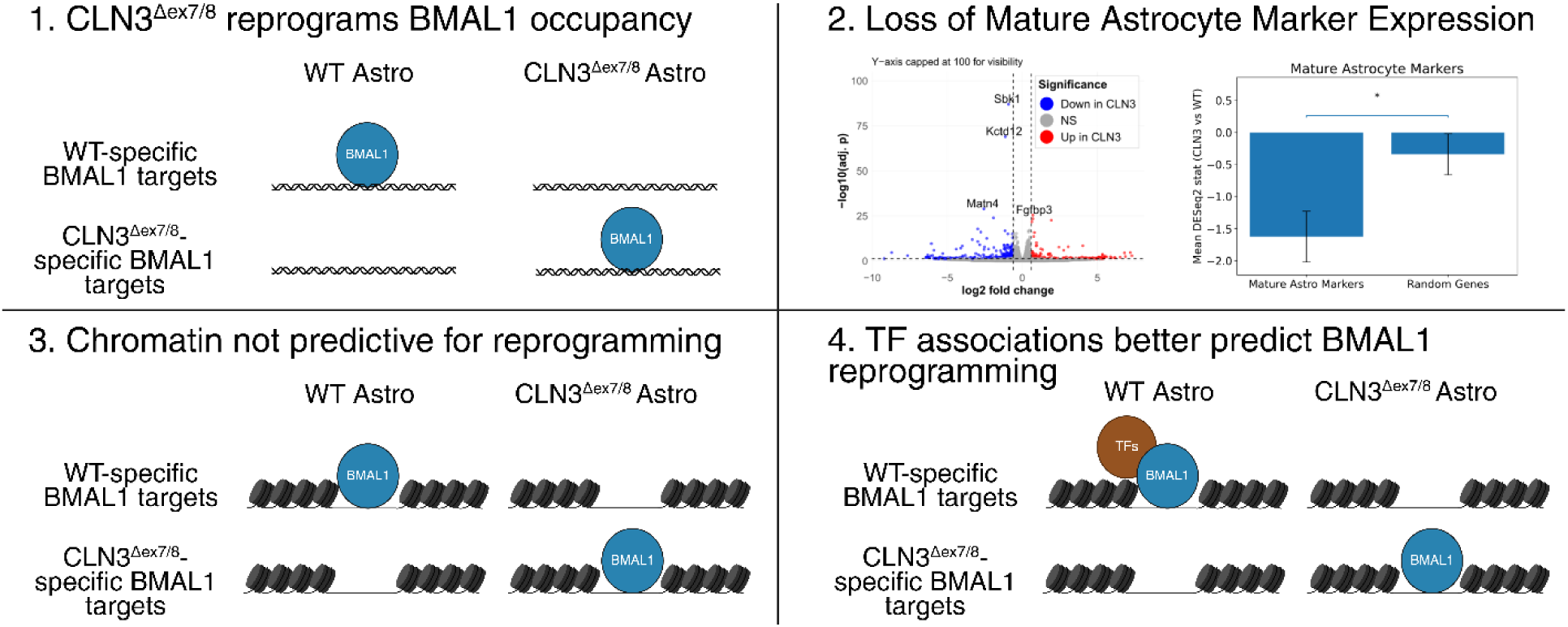

## INTRODUCTION

Astrocytes regulate neurotransmitter recycling, metabolic and redox support, ion and water homeostasis, and synapse formation and pruning, processes that shape neuronal survival and circuit function across the lifespan (1). Disruption of these astrocytic functions is increasingly recognized as a key contributor to neurodegenerative disease progression (2). Circadian rhythms have also recently emerged as a potential modifier of neurodegeneration (3), as many core astrocyte homeostatic functions are regulated by the circadian clock. Disruption of astrocyte homeostatic programs is associated with reactive astrogliosis, a ubiquitous feature of CNS disorders that exhibit neurodegeneration (4, 5), characterized by a gain of neurotoxic, pro-inflammatory signaling that can drive excitotoxic stress and directly promote neuronal death (6). Disruption of circadian rhythms also induces reactive astrogliosis, suggesting that circadian regulation of astrocyte function may be compromised during neurodegeneration (7).

A recent study provides evidence that the astrocytic circadian clock is disrupted at the level of transcriptional regulation: in an Alzheimer’s disease model, astrocyte-specific circadian programs are reprogrammed, with metabolic pathways losing rhythmicity while PI3K and TNF signaling pathways gain rhythmic oscillation (8). These findings suggest that neurodegeneration is associated not simply with dampened circadian output, but with a redistribution of circadian regulatory control across astrocyte gene programs. This general phenomenon of circadian reprogramming has been observed in diverse contexts including across tissues (9), as well as within various diet or disease models (10–12), so it would not be surprising for circadian gene expression to be reprogrammed during neurodegeneration as well.

Circadian gene expression is governed by a cell-autonomous molecular clock. At the core of this clock, the CLOCK:BMAL1 heterodimer binds E-box elements to activate transcription of Per and Cry genes, whose protein products provide inhibitory feedback on CLOCK:BMAL1 activity, generating ∼24-hour rhythms (13). Among core clock transcription factors (TFs), BMAL1 is uniquely nonredundant, as its deletion abolishes circadian behavioral rhythms and markedly blunts rhythmic gene expression (14). Astrocyte-specific deletion of BMAL1 is sufficient to induce a reactive astrocyte state, suggesting that astrocyte reactivity in neurodegenerative disease may reflect impaired BMAL1 function (7). Consistent with its role in maintaining astrocyte homeostasis, BMAL1 has been proposed as a mechanistic link between circadian disruption and neurodegeneration (15, 16).

Several previous studies have examined BMAL1 expression patterns in neurodegenerative disease patients (17), but the results have been inconsistent, with some reporting loss of rhythmic BMAL1 expression and others finding preserved but phase-shifted rhythms. Moreover, prior work has emphasized that a TF’s mRNA abundance is not a reliable proxy for its regulatory activity (18). Instead, we have focused on whether BMAL1-DNA binding is altered in the setting of neurodegenerative pathology, rather than relying on disruption of its mRNA expression alone. To date, BMAL1 genomic occupancy has not been profiled *in vivo* in astrocytes, or in a neurodegenerative context in general. Direct measurement of BMAL1 binding is therefore required to identify its direct gene targets and to mechanistically assess its regulatory function. We seek to develop new technologies to record BMAL1 binding specifically within astrocytes.

Given evidence that BMAL1 can shape astrocyte activation programs, we asked whether BMAL1 occupancy is reprogrammed in a neurodegenerative setting where astrocyte state changes are an early and informative feature of pathology, and may contribute to subsequent neuron loss. We focused on neuronal ceroid lipofuscinosis (NCL), a group of fatal inherited disorders of childhood that display profound neurodegeneration as a result of lysosomal dysfunction (19). Mouse and large animal models of NCL consistently display localized astrocytic and microglial activation that emerges early in disease progression and predicts the later pattern of selective neuron loss. Amongst the NCL subtypes we chose CLN3 disease, which progresses more slowly than earlier onset forms such as CLN1 and CLN2 disease, providing a more prolonged but defined window to interrogate astrocyte dysregulation before subsequent widespread neurodegeneration (20). CLN3 deficiency produces functional abnormalities in primary astrocytes, such as blunted activation, abnormal morphological responses, and altered secreted factors (including cytokines/chemokines and neuroprotective proteins), impaired calcium signaling, and disrupted glutamate handling, which reflect early glial dysfunction (21). We performed experiments during the first two postnatal months to capture early, mechanistically informative BMAL1 regulatory changes before late-stage secondary cascades occur (reactive glial responses, circuit disruption, shifting cell-state composition) (22, 23). *Cln3^Δex7/8^*mice bear the mouse equivalent 1.02kb deletion in *CLN3* that is present in ∼85% of human CLN3 disease-causing alleles (24, 25). These *Cln3^Δex7/8^* mice also show altered circadian-related behavior at later ages, suggesting that BMAL1 dysregulation is plausible within this widely used model of CLN3 disease (22).

To determine if BMAL1 function is altered within *Cln3^Δex7/8^* astrocytes, it is first critical to identify the genomic binding sites of BMAL1 in astrocytes *in vivo*. However, this is challenging because the brain is a complex tissue and conventional genome-wide TF occupancy assays are difficult to restrict to a single cell type at sufficient scale. We therefore developed MACS-Calling Cards (MACS-CC), which combines *in vivo* piggyBac Calling Cards with magnetic-activated cell sorting (MACS) to enrich astrocytes prior to library preparation, enabling astrocyte-enriched mapping of BMAL1 genomic occupancy. Using this method, we were successfully able to map *in vivo* BMAL1 occupancy and, when applying MACS-CC to wild-type and *Cln3^Δex7/8^* mice, observed extensive BMAL1 reprogramming. Wild type (WT)-specific sites preferentially map near glial differentiation-associated gene programs, whereas CLN3 disease-specific sites lack coherent functional enrichment, consistent with a loss of WT-like functional targeting in CLN3 disease. We next tested whether chromatin accessibility remodeling could explain BMAL1 retargeting. If accessibility were a primary driver, gained BMAL1 sites would be accompanied by increased accessibility. Instead, promoter accessibility is broadly remodeled in CLN3 disease, while accessibility changes at BMAL1 peaks and their correlation with binding changes are minimal, arguing against an accessibility-driven model. Finally, motif occupancy analyses reveal disruption of cofactor-associated motif architecture near BMAL1 motifs, supporting a model in which altered cooperative partner associations contribute to BMAL1 retargeting in the early stages of a neurodegenerative cascade. Our findings provide new insights into astrocytes in CLN3 disease, suggesting impaired development. Overall, we developed a generalizable strategy for cell type-specific TF occupancy mapping *in vivo* and revealed early rewiring of astrocyte circadian regulatory control in neurodegenerative disease.

## MATERIAL AND METHODS

### Cell Culture

HepG2 cells were maintained in DMEM+10% FBS, 1x Penstrep in 6 well plates. Calling card constructs (with puro as the marker gene) transfected through neon electroporation (2*10^6 cells/well). Puro selection performed with 3ug/mL puro. Puro selection was done over 5 days to allow for cells without insertions to die off, a non-transfected well was used to determine that all cells without insertions had died at that point. Plasmids used for CC experiments *in vitro* are detailed in Supplementary Table S1.

### Mouse Husbandry

All animal procedures were performed in accordance with NIH guidelines and were approved by the Washington University School of Medicine Institutional Animal Care and Use Committee (IACUC). Mice were housed at Washington University in St. Louis under standard vivarium conditions on a 12:12-hour light:dark cycle with food and water available ad libitum. The Aldh1l1-TRAP line was obtained from the laboratory of Joseph D. Dougherty, and *Cln3^Δex7/8^* has been utilized as a genetically accurate model of CLN3 disease for two decades (24) and were maintained on a C57BL/6J strain background. Wild-type controls were C57BL/6J mice (The Jackson Laboratory, stock #000664) and were maintained in the same facility. The *Cln3^Δex7/8^* model line was maintained as homozygotes; the Aldh1l1-TRAP line was maintained as heterozygotes with littermates acting as controls for MACS and flow cytometry. Genotyping of GFP was confirmed by PCR using DNA from ear biopsies extracted with the PCRBIO Rapid Extract Lysis Kit (PB15.11-08). Genotyping primers are listed in Supplementary Table S2.

### AAV injection

Viruses were packaged by the Hope Center Viral Vectors Core. AAV injections were performed as described in (26). Viral titers (viral genomes [vg] per milliliter) were ∼1.0 × 10. Neonatal mice (P0-P1) received intracerebroventricular (ICV) injections of Calling Cards AAVs encoding the piggyBac transposase (unfused or BMAL1 fusion) and donor/self-reporting transposon. AAV stocks were mixed 1:1 immediately prior to injection (6 µL total per pup). Pups were anesthetized by hypothermia on ice (5-8 min; ≤15 min maximum) and injected bilaterally using a Hamilton syringe fitted with a depth guard to ensure a consistent 2 mm insertion depth. Virus was delivered at 1 µL per site at three sites per hemisphere (1 µL/s); after each injection, the needle was held in place for 5 s before withdrawal to minimize backflow. Following injection, pups were placed on a low warming pad for recovery (5-10 min) and returned to the home cage with the dam. Plasmids used for CC experiments *in vivo* are detailed in Supplementary Table S3.

### Cloning

Constructs were assembled using Gibson Assembly (NEB E2611S). Several parent plasmids were existing Mitra lab or collaborator lab reagents that have not been formally published; annotated plasmid maps and/or complete sequences can be provided upon request.

*pRM2097 (CMV_mBmal1_hyPBase; in vitro BMAL1-hyPBase):* pRM2097 was generated by replacing the Brd4 coding sequence in a Brd4-hyPBase Calling Cards fusion plasmid (pRM1878) with mouse *Bmal1* cDNA (Sino Biological MG57171-U). The resulting construct expresses a mouse BMAL1-hyPBase fusion from a CMV promoter for cell culture Calling Cards experiments.

*pRM2095 (AAV_CAG_mBMAL1_hyPBase; in vivo AAV BMAL1-hyPBase):* pRM2095 is an AAV transfer plasmid expressing mouse BMAL1–hyPBase from a CAG promoter. It was generated using an AAV backbone plasmid obtained from the Dougherty lab (PGK-TetO3G AAV transfer vector) by replacing the entire cassette between the AAV ITRs with a CAG promoter + BMAL1–hyPBase expression cassette derived from pRM2097.

*pRM2096 (AAV_CAG_hyPBase; in vivo AAV unfused hyPBase):* pRM2096 was derived from pRM2095 design by substituting the BMAL1-hyPBase fusion ORF with an unfused hyPBase ORF sourced from pRM1114, while keeping the same AAV transfer backbone and CAG promoter configuration. This construct expresses hyPBase alone for unfused transposase Calling Cards controls *in vivo*.

All final constructs were validated b*y* restriction enzyme digestion and sequencing of the entire plasmid through Plasmidsaurus.

### Astrocyte enrichment

Astrocytes were enriched using a protocol adapted from (27), with modifications to specifically sort for astrocytes. Briefly, mouse cortices were dissected and then dissociated according Miltenyi’s protocol for the Adult Brain Dissociation Kit (130-107-677). Once the cells were in a single cell suspension, myelin was removed using Miltenyi’s Myelin Removal Beads Kit II (130-096-733) according to the manufacturer’s instructions, and the flow-through was retained. Dead cells were subsequently removed using Miltenyi’s Dead Cell Removal Kit (130-090-101) following the manufacturer’s protocol, and again the flow-through was collected. Finally, astrocytes were isolated using Miltenyi’s ACSA2 Isolation Kit (130-097-678) according to the manufacturer’s instructions, with the ACSA2-bound fraction considered to be astrocytes. For Calling Cards samples, astrocyte enrichment was performed at 2 months of age; for RNA-seq and MNase-seq samples, enrichment was performed on equal numbers of 1-month-old and 2-month-old mice, to match the time period that Calling Cards records binding.

### Reference annotations and gene assignment for computational analyses

Transcription start sites (TSSs) were derived from the *Gencode vM25 basic* mouse annotation GTF. Transcript features were filtered to protein-coding transcripts with APPRIS principal isoform tags 1–2. Promoter windows were then defined as ±2 kb around each TS. “Closest” genes were identified using HOMER’s annotatePeaks.pl. For ShinyGo gene-set enrichment, a nearby gene was defined as the closest gene within 5kb of the TSS, according to HOMER’s distance to TSS metric. ShinyGo version is 0.85.

### bigWig Normalization

To enable direct comparison between two condition-averaged bigWig signal tracks, we performed an additional post-hoc signal normalization for all bigwig comparisons (either CC, MNase-seq, ATAC-seq, or FAIRE-seq). First, a common set of reference regions was defined for estimating the between-track normalization function. Transcription start site (TSS) annotations were expanded to ±1 kb in a strand-aware manner using bedtools slop to generate promoter windows. The two bigWig tracks were then summarized over a shared reference BED of sites using deepTools multiBigwigSummary BED-file, producing a matrix of per-region signal values.

We next applied a symmetric quantile-normalization strategy so that both tracks were mapped to the same intermediate (“midpoint”) target distribution, minimizing bias toward either condition. Using the summarized per-region values, a custom script fit two monotonic quantile mapping functions (one for each input track) to a midpoint distribution defined as the mean of the two empirical quantile curves. The mapping functions were parameterized using spline knots and written as tab-delimited lookup tables. These fitted mappings were then applied genome-wide to the full-resolution bigWig files, replacing each original signal value with its mapped midpoint-normalized value; negative values produced by the remapping were clipped to zero. If needed, the resulting quantile-normalized bigWigs were used as inputs to deepTools bigwigCompare to generate a difference track using a log2 operation.

### MNase-seq

#### MNase digestion

MNase digestion was performed immediately following astrocyte isolation. Cells were fixed for 10 minutes in 1% formaldehyde and the reaction was quenched for 5 minutes with 2.5 M glycine, added to a final concentration of 0.125 M. The cells were then washed twice in 1x PBS. Nuclei were prepared by adding 100 μl of Nuclei Prep Buffer (Zymo EZ Nucleosome DNA Prep Kit, D5220) per 1 million cells. After centrifugation, the cells were resuspended in 1x Takara MNase Digestion Buffer. Chromatin was digested with 4 units of Takara MNase (2910A) per 1,000,000 cells. The reaction was stopped by adding 5x MN stop buffer from the EZ kit. Subsequently, 4.8 μl of 5 M NaCl and 2 μl of proteinase K were added per 1 million cells, and samples were incubated at 65°C overnight. The following day, DNA was isolated using the EZ kit according to the manufacturer’s instructions.

#### MNase-seq library prep and sequencing

MNase-seq libraries were prepared using the NEBNext Ultra end repair and A-tailing module (NEBNext End Prep, E7442), followed by adapter ligation using the NEBNext Ultra Ligation Module with Quick Ligation (E7445). Libraries were purified using Agencourt AMPure XP beads (Beckman Coulter, A63880). Adapter-ligated DNA was PCR-amplified using Phusion High-Fidelity DNA Polymerase with NEB primers (NEB, E6440S), followed by a second cleanup with Agencourt AMPure XP beads. All samples were then submitted to the the Washington University Genome Technology Access Center (GTAC) to be deeply sequenced with the Illumina NovaSeq X Plus. MNase-seq primer and adapter sequences are listed in Supplementary Table S4.

#### MNase-seq data processing

Paired-end MNase-seq FASTQ files were aligned to the mm10 reference genome using Bowtie2 v2.3.5. Alignments were performed in paired-end mode with an expected insert-size range of 110-210 bp (-I 110 -X 210), requiring properly paired reads, and reads were trimmed to 40 bp prior to alignment. SAM files were converted to BAM and unmapped reads were removed, followed by coordinate sorting and indexing. To restrict analyses to canonical chromosomes, BAMs were filtered to chr1-chr19, chrX, chrY, name-sorted, and processed with samtools fixmate. BAMs were then coordinate-sorted and PCR duplicates were removed with samtools markdup -r.

To generate genome-browser tracks, processed replicate BAMs for each condition were supplied to DANPOS3 for differential analysis using the standard danpos.py. DANPOS3 outputs included pooled per-condition WIG tracks (normalized/smoothed signal) and a differential WIG track (Poisson-based difference). Finally, WIG files were converted to bigWig format using the UCSC wigToBigWig utility. This produced bigWig tracks for each condition’s normalized/smoothed MNase signal and for the DANPOS differential signal.

#### Differential promoter NFR calling from MNase-seq

Construction of strand-aware NFR and flanking windows. For each TSS, three windows were constructed: (i) an NFR window centered on the TSS (−75 to +75 bp), (ii) a left flank (−300 to −150 bp), and (iii) a right flank (+150 to +300 bp). All windows were strand-aware (i.e., “left” and “right” are defined relative to transcript orientation), clipped to chromosome bounds, and deduplicated.

MNase fragment counting with featureCounts. MNase-seq read pairs from all BAMs were quantified over the NFR/flank windows using featureCounts with SAF annotations (1-based start), requiring properly paired reads, minimum mapping quality MAPQ≥10, and restricting counted fragments to a mononucleosome-sized insert window of 120–200 bp.

Per-gene “open promoter NFR” QC metric. Counts were aggregated across all CLN3 samples and all WT samples separately by summing the featureCounts columns corresponding to each directory, and library sizes were defined as the total summed counts across all windows per group.

For each gene, group-level counts for NFR, left flank, and right flank were computed, and the flank mean was defined as the average of left and right flank counts. The NFR depletion ratio was computed as: Ratio = FlankMean / NFR (with defensive handling for zero NFR counts). FlankMean was converted to CPM using the group library size. A promoter NFR was considered PASS (i.e., “open”) within a group if it satisfied both thresholds: FlankMean_CPM ≥ 0.5 and FlankMean/NFR ≥ 1.3 This produced group-specific PASS calls and BED files of PASS NFRs for CLN3 and WT.

Differential NFR set definitions (presence/absence). “Differential NFRs” were defined by overlap relationships between the CLN3-pass and WT-pass NFR BEDs using strand-aware bedtools operations: CLN3-specific = present in CLN3-pass and not overlapping WT-pass; WT-specific = present in WT-pass and not overlapping CLN3-pass; Both = present in both; Neither = NFRs that overlap neither pass set.

### Calling Cards

#### Calling Cards Library Prep and Sequencing

Total RNA extraction after MACS was done using the Qiagen RNeasy Plus Mini Kit (Cat no. 74134) according to the manufacturer’s protocol. All of the total RNA was used for reverse transcription. First-strand cDNA was generated by oligo(dT) priming with the SMART_dT18VN primer and Maxima H Minus reverse transcriptase in the presence of RNase inhibitor, followed by heat inactivation and RNase H treatment to remove the RNA strand, yielding cDNA for downstream library construction. Self-reporting transposon (SRT) transcripts were then PCR-amplified from cDNA using KAPA HiFi HotStart with a reporter specific primer (tdtomato or puromycin) and SMART reverse primers (cycle number minimized/optimized to avoid over-amplification; typical program uses ∼20 cycles with 98°C denaturation, 65°C annealing, and long 72°C extension), and amplicons were size-selected/cleaned with a 0.6× AMPure XP bead cleanup and QC’d by Qubit. Purified SRT PCR products were converted into sequencing-ready libraries by Nextera XT tagmentation (ATM + TD buffer at 55°C), neutralization, and an indexing PCR using a barcoded piggyBac (OM-PB) primer together with a Nextera N7 index primer to enable multiplexing, followed by a final 0.6× bead cleanup and fragment distribution QC prior to pooling and Illumina sequencing. A fraction of the samples were checked for sequencing quality using WashU’s DNA Sequencing Innovation Lab (DSIL)’s spike in program with the Illumina MiSeq or Illumina Miniseq. All samples were then submitted to the the Washington University Genome Technology Access Center (GTAC) to be deeply sequenced with the Illumina NovaSeq X Plus. All sequencing data was combined at the end for each sample. Calling Cards primer sequences are listed in Supplementary Table S5.

#### Calling Cards Insertion Calling

Calling Cards FASTQ reads were processed with a custom bash pipeline to generate per-sample insertion files for peak calling. Briefly, reads were first adapter-trimmed and demultiplexed by requiring an exact 5′ match to the OM-piggyBac transposon sequence using cutadapt with zero mismatches/indels and discarding untrimmed reads, then SRT barcodes were extracted with umi_tools extract, followed by an additional exact-match trim of the transposon sequence and trimming of trailing Nextera adapters (cutadapt). Trimmed reads were aligned to the reference genome with Bowtie2 and converted to BAM; mapped reads were retained, then sorted and indexed. Transposon insertion sites were then computed per read by assigning the insertion point to the alignment start (forward reads) or end (reverse reads), correcting for soft clipping, and encoding the genomic interval immediately preceding the insertion. For piggyBac, reads were optionally filtered to those matching the expected “TTAA” insertion motif. Finally, unique insertions were aggregated into an insertion table (“qbed” file) by counting identical insertion sites; these insertion files were then used as input for downstream peak calling.

#### Differential Calling Cards Peak Calling

Differential Calling Cards peak sets were generated using a custom pipeline that (i) defines a unified candidate peak set and (ii) classifies those peaks as condition-enriched (CLN3-specific/WT-specific) or shared. For each condition, peaks were called with PyCallingCards using the MACCs (“cctools”) framework with the parameters pvalue_cutoffbg=0.05, window_size=800, step_size=400, pvalue_cutoffTTAA=0.005, lam_win_size=1,000,000, pseudocounts=0.4. Peaks from the two TF conditions were then concatenated and collapsed into a nonredundant “union” peak set by sorting and merging intervals with bedtools merge using a 1 kb merge distance (-d 1000). To call differential peaks, we quantified insertions from the two TF qbed files across the union peak set using pycallingcards.pp.make_Anndata, then tested TF_C1 vs TF_C2 per peak with Fisher’s exact test and ranked peaks by multiple-testing–adjusted p-values. Peaks with adjusted p < 0.05 enriched in TF_C1 were designated CLN3-specific peaks, and peaks enriched in TF_C2 were designated WT-specific peaks (written as separate BED files). Finally, “shared” peaks were defined as the remaining union peaks not assigned to either differential class, implemented with bedtools intersect -v against the condition-specific BED files.

#### Calling Cards bigWig generation

Genome-wide insertion signal tracks were generated from Calling Cards .qbed files using a shell pipeline that bins insertion intervals, normalizes counts, smooths the signal, and converts the result to bigWig format. For each .qbed file, fixed-width genome bins were first created with bedtools makewindows for a bin width of 50bp. The number of insertion events overlapping each genomic bin (≥1 bp overlap) was then computed with bedtools coverage -counts. To account for library size, bin counts were converted to counts-per-million insertions (CPM). Total insertions were computed as the number of non-header lines in the .qbed, and a scaling factor of 10^6/total insertions was applied to the bedGraph count column to produce a CPM-normalized bedGraph. CPM signal was smoothed using a centered rolling mean (“boxcar”) across W adjacent bins (default W=7, corresponding to ∼350 bp when BIN=50). Smoothing was performed separately within each chromosome: for each bin, the mean CPM across a window of ±⌊W/2⌋ bins was computed, with window boundaries clamped at chromosome edges. The smoothed bedGraph was sorted by chromosome and start coordinate and converted to bigWig using UCSC’s bedGraphToBigWig.

#### Motif occupancy

To quantify motif occupancy around condition-specific NFRs, we expanded the NFRs to 2kb around the TSS and labeled these condition-specific promoters. So a WT-specific NFR would become a WT-specific promoter, for example. We utilized CLN3-specific, WT-specific, and shared promoters for the rest of quantifying motif occupancy. We utilized BMAL1 peaks with their original sizes.

#### Motif instance calling

For each peak/promoter set compared, the script first created a merged union of intervals and then identified occurrences of a BMAL1 control motif across the union using HOMER annotatePeaks.pl. For each HOMER motif file, annotatePeaks.pl was run on the peaks/promoters set and the shared set to export motif-instance BED files for each motif.

#### Occupancy metrics and hypothesis tests

##### Motif occupancy within promoters

To calculate general occupancy at promoters, we summarized the motif instances of each promoter set into a *promoter-level occupancy* comparison, i.e. for each promoter set, it collapses “how many motif instances exist” into whether each promoter is motif-positive or motif-negative (motif-positive = promoter overlaps ≥1 motif instance). It then builds a 2×2 contingency table per motif and computes Fisher’s exact test p-values for difference in occupancy between promoter sets.

##### Motif occupancy near BMAL1 motifs within BMAL1 peaks

We performed a similar analysis to quantify *motif occupancy nearby BMAL1 motifs within BMAL1 peaks*: For each BMAL1 peak set, we defined windows of ±200 bp around each BMAL1 control-motif instance. For each motif-of-interest, we scored each BMAL1-centered window as motif-present if it contains ≥1 motif instance. We then compared the fraction of BMAL1-windows that are motif-present between WT/CLN3-specific BMAL1 peaks and shared BMAL1 peaks via Fisher’s exact test.

##### Calling highlighted motifs

We applied multiple-testing correction using Benjamini–Hochberg FDR to Fisher’s exact test p-values in all occupancy tables. Motifs were counted as “highlighted” (and subsequently highlighted in the resulting scatterplot) if they met both the FDR and effect cutoffs for both the *promoter-level occupancy* and *motif occupancy nearby BMAL1 motifs within BMAL1 peaks*.

### RNA-seq

Total RNA extraction after MACS was done using the Qiagen RNeasy Plus Mini Kit (Cat no. 74134) according to the manufacturer’s protocol. RNA-seq libraries were prepared in-house from total RNA using the NEBNext UltraExpress® RNA Library Prep Kit (New England Biolabs) according to the manufacturer’s instructions. Indexes used from NEBNext Multiplex Oligos for Illumina® (E6440S). Final libraries were quantified, pooled, and submitted to the Washington University Genome Technology Access Center (GTAC) for sequencing. GTAC performed sequencing on the Illumina NovaSeq X Plus (paired end reads extending 150 bases) and generated gene-level RNA-seq count tables from the resulting sequencing data using their standard RNA-seq processing workflow; these GTAC-provided count tables were used to assemble the final count matrix for downstream analyses.

#### RNA-seq computational analysis

##### Removing contamination from non-astrocytes

Because around 20% of cells in ACSA2-bound fraction are potentially non-astrocytes (Figure 2B), we wanted to address the possibility that CLN3 and WT mice could differ in the cell makeup of the contamination, in that certain cell types could be present in different abundances, skewing what genes are called as DEGs. Because we are not studying non-astrocytes, we just wanted to remove these differences from affecting our analysis of astrocytes. To assess potential non-astrocyte contamination in the bulk astrocyte RNA-seq dataset, we quantified sample-level “enrichment scores” for curated marker panels representing major CNS cell types using a custom Python workflow. For each potential cell type contaminant, the enrichment score for a given sample was calculated as a mean z-score across a set marker genes for that cell type. To visualize whether marker enrichment differed systematically by genotype/condition, the script generated scatterplots of Astro score (x-axis) versus each non-astro panel score (y-axis). Based on inspection of these scatterplots, if a given non-astro panel showed an apparent condition-specific shift (in this dataset, this was observed for microglia and oligodendrocyte marker panels), the corresponding enrichment scores were added as numeric columns to the sample metadata table and treated as covariates in the differential expression model.

#### Differential expression testing

Differentially expressed genes (DEGs) between CLN3 and WT astrocyte samples were identified using a custom R pipeline built around DESeq2 with surrogate variable analysis (SVA) to reduce unmodeled technical/latent variation. The script reads the same filtered count matrix and metadata table and sets WT as the reference condition and CLN3 as the treated condition. The count table was parsed by detecting a gene identifier column (preferring Feature when present), removing duplicate gene rows, dropping common annotation columns, and converting the remaining sample columns into an integer count matrix (NA values set to 0 and counts rounded to integers). Metadata were required to contain sample and condition columns; samples not present in the count matrix were dropped, and the count matrix was reordered to match the metadata sample order. The condition factor was releveled to ensure WT was the reference level, and the script verified that the treated level (CLN3) was present.

To incorporate contamination/marker enrichment covariates (and any other relevant sample-level variables), the script automatically identified additional covariate columns in the metadata beyond sample and condition that showed variability across samples; character covariates were coerced to factors. These covariates were combined with condition to form the initial design formula. Genes were prefiltered by requiring a minimum total count across all samples (prefilter_min_total = 10). Surrogate variables were then estimated using svaseq on DESeq2 size-factor–normalized counts, using a full model that included condition plus all varying covariates and a null model that included only the covariates (excluding condition). DESeq2 was run using glmGamPoi as the fitting engine and a likelihood ratio test (LRT) to test the overall effect of condition while controlling for surrogate variables and included covariates; the reduced model dropped condition but retained surrogate variables and covariates. In parallel, the script performed a Wald test for the CLN3 vs WT coefficient and applied an effect-size threshold using lfcThreshold = 0.2 with altHypothesis = “greaterAbs” (i.e., testing whether the absolute log2 fold-change exceeds the specified threshold), then ordered results by adjusted p-value. Log2 fold-changes were additionally shrunken using apeglm for effect-size estimation while retaining the thresholded Wald test p-values for significance decisions. For final DEG reporting, we applied an additional, more stringent post hoc cutoff to the Wald results, requiring an absolute log2 fold-change > 0.6 and adjusted p-value (FDR) < 0.05; genes meeting both criteria were reported as DEGs.

### ChIP-seq, ATAC-seq, and FAIRE-seq re-analysis

Raw sequencing runs were downloaded from NCBI SRA and converted to FASTQ using the SRA Toolkit (SRA “prefetch” followed by FASTQ extraction with paired-end splitting when applicable). FASTQ files were gzip-compressed for storage. Reads were aligned to mm10 using Bowtie2 in local alignment mode. For ATAC-seq and FAIRE-seq datasets, paired-end reads were aligned using a “very-sensitive” local preset and a maximum fragment length of 2000 bp. For ChIP-seq datasets, reads were aligned as single-end in local mode. After alignment, unmapped reads were removed, alignments were coordinate-sorted and indexed, and BAMs were restricted to canonical chromosomes (autosomes, X/Y, and mitochondrial) prior to downstream analysis.

#### Differential ChIP-seq peak calling

For each ChIP-seq replicate, peaks were called against the corresponding input control (either a pooled input or replicate-matched inputs) using MACS3 with summit calling enabled. Peaks were called using a stringent q-value threshold of 1×10⁻⁵, and peak summits were used as the anchor for subsequent quantification.

To enable differential testing on a common set of genomic intervals, peak-centered windows were constructed around called summits for each sample and then merged across all samples to form a union set of candidate regions. In this workflow, 500-bp windows centered on summits (±250 bp) were used, and overlapping windows were merged. Read counts for each replicate within each merged region were computed using bedtools multicov, requiring a minimum mapping quality of 30.

In parallel, an “occupancy” matrix was generated to indicate whether each region was “present” in each replicate’s peak set. Occupancy was defined using a midpoint rule: for each merged region, a 1-bp midpoint interval was intersected with that replicate’s peak calls, and regions whose midpoint overlapped at least one peak were marked occupied (1), otherwise unoccupied (0). These per-region count and occupancy matrices were assembled into a single table for downstream statistical modeling.

Differential binding between conditions was tested using a limma-voom workflow on the region-by-sample count matrix. Counts were TMM-normalized, transformed with voom, fit with a linear model including condition, and moderated using empirical Bayes shrinkage; log2 fold-changes were computed as Condition 2 vs Condition 1, and p-values were adjusted using the Benjamini–Hochberg procedure. Regions were considered differentially bound at BH-adjusted q-value < 0.20, and were classified as Condition 1-enriched vs Condition 2–enriched based on the sign of the fitted log2 fold-change (negative = higher in Condition 1; positive = higher in Condition 2).

#### ATAC-seq and FAIRE-seq bigWig Generation

For ATAC-seq datasets, BAMs were additionally filtered to retain high-confidence alignments (MAPQ ≥ 30), remove unmapped reads and marked duplicates, and then Tn5-shifted using an ATAC-specific shifting procedure prior to downstream use. Shifted BAMs were coordinate-sorted and indexed (FAIRE-seq datasets did not undergo ATAC shifting in this pipeline.)

To generate comparable genome-wide coverage tracks, per-sample TMM scaling factors were computed from the region count matrix using edgeR: effective library sizes were computed as lib.size × norm.factor, a target depth was set to the geometric mean of effective library sizes, and each sample’s BigWig scale factor was defined as target / effective_library_size.

For each ATAC-seq or FAIRE-seq replicate BAM, normalized coverage BigWigs were created using bamCoverage with 25-bp bins, applying the precomputed TMM scaleFactor (with no additional built-in normalization), ignoring duplicates, requiring MAPQ ≥ 30, and excluding blacklisted genomic regions (mm10 blacklist).

Finally, replicate BigWigs were averaged within each condition to produce a single mean track per condition using weighted addition (equal weights of 1/n per replicate) implemented via bigwigCompare.

### Circadian rhythmicity (RAIN) re-analysis

Gene-level circadian rhythmicity was re-analyzed from a gene-by-time CPM matrix (8) using the RAIN algorithm. Expression values were imported from a CSV file in which the first column contained gene identifiers and subsequent columns corresponded to timepoints (with replicate samples encoded as paired columns per timepoint, e.g., X0 and X0.1 for two replicates). Columns were ordered by circadian time and replicate index so that replicates remained adjacent within each timepoint prior to analysis.

RAIN was run on the transposed expression matrix (timepoints as rows, genes as columns) using a 2-hour sampling interval (Δt = 2 h), a 24-hour period, and two replicate series per timepoint (nr.series = 2). Peak detection boundaries were constrained to the middle portion of the cycle (peak.border = 0.3–0.7), using the “independent” method, and adjusted p-values were computed with the adaptive Benjamini–Hochberg procedure (adjp.method = “ABH”). For each gene, RAIN output statistics (including raw p-values, adjusted p-values, inferred phase, peak-shape, and period) were collected.

### Phase-normalized heatmaps for near-peak and mid-cycle expression

Heatmaps were generated from the same expression matrix as RAIN analysis by extracting timepoint columns and (optionally) collapsing replicate columns (taking the mean) belonging to the same integer timepoint into a single value per time (e.g., averaging X0 and X0.1), producing a genes-by-time matrix for each gene set. Expression values were optionally transformed (log2(CPM+1)) and z-scored across time within each gene to emphasize rhythmic shape over absolute level.

For the phase-normalized representation, each gene’s timecourse was “peak-aligned” by circularly shifting its row so that the timepoint of maximal expression occurred in the first column. Each peak-aligned gene row was summarized into two values: the mean expression within the near-peak “hours-since-peak” window (22hrs (aka -2hrs), 0hrs, 2hrs since peak) and the mean expression within the user-defined mid-cycle window (10hrs, 12hrs, and 14hrs since peak). Genes were ordered using the metric “windowdiff”, defined as mean(near-peak window) − mean(mid-cycle window), with near-peak and mid-cycle windows specified as hours since peak).

To make groups comparable, each phase-normalized gene set was downsampled to the size of the smallest set (sampling rows without replacement; random sampling uses a fixed seed and preserves the original ranked order), and then an optional plotting cap was applied so only the first 150 rows were displayed after downsampling.

Finally, the two means of each gene (near-peak and mid-cycle) were organized into a per-gene 2-column matrix (near-peak, mid-cycle) for each gene set and plotted as a side-by-side panel across groups with a shared color scale.

## RESULTS

### The Calling Cards technology accurately records BMAL1 binding *in vitro*

To test whether the early stages of a neurodegenerative cascade reprogram BMAL1’s binding patterns in *Cln3^Δex7/8^* astrocytes, we sought to develop a method to map BMAL1 binding sites *in vivo* with astrocyte specificity - an analysis that is difficult in brain tissue using conventional genome-wide occupancy assays. To overcome this, we combined Magnetic-Activated Cell Sorting with Calling Cards (MACS-CC): MACS is an immunomagnetic enrichment method in which cells labeled with antibody-conjugated magnetic microbeads are retained in a magnetic field (often a column) while unlabeled cells flow through, enabling rapid enrichment of a target population from brain dissociates (28). Calling Cards (CC) is a method to record TF-DNA binding in which a TF is coupled to a transposase so that TF-DNA interactions are converted into nearby transposon insertions, creating a stable genomic “record” of binding events that can be recovered later by sequencing, and can be performed with viral delivery strategies suitable for mammalian tissues (26, 29–32). CC allows recording of BMAL1 binding before the perturbations of dissociation and sorting, which are known to induce *ex vivo* cellular stress programs and can confound downstream molecular readouts (33, 34).

Accordingly, before introducing astrocyte enrichment, we first validated that BMAL1 CC works on its own in a cell-based system: i.e., that BMAL1-directed insertions are robustly recoverable and show the expected enrichment patterns, providing the foundation for the subsequent MACS-CC astrocyte-specific experiments. We performed BMAL1 CC in HepG2 cells, where BMAL1 ChIP-seq data is available for benchmarking (Encode ENCSR794LVK). We fused BMAL1 to the hyperactive piggyBac transposase (hyPBase). In the presence of the fusion protein, hyPBase inserts a self-reporting transposon (SRT) near BMAL1-bound DNA (Figure 1A). The SRT subsequently produces RNA, and we recover insertions through a modified bulk RNA-seq protocol (termed CC library prep). Each experiment included an unfused hyPBase control to measure the baseline, BMAL1-independent transposon insertion pattern, so BMAL1-fused CC insertions can be interpreted as enrichment above that background. In HepG2 cells, BMAL1-directed CC insertion pileups, when compared to the unfused background, produced BMAL1 CC peaks that overlapped BMAL1 ChIP-seq peaks in a genome-browser view, specifically at the core clock target *Dbp*, which BMAL1 is known to bind to (13) (Figure 1B). Genome-wide, BMAL1 ChIP-seq signal centered on BMAL1 CC peak centers, indicating that insertion peaks faithfully report BMAL1 occupancy (Figure 1C). Consistent with expected sequence specificity, the canonical E-box motif was the top enriched known motif in BMAL1 CC peaks (Figure 1D). Finally, genes near BMAL1 CC peaks were enriched for circadian rhythm pathways by ShinyGO (Figure 1E). Together, these results indicate that BMAL1 Calling Cards accurately captures BMAL1 binding and can serve as a practical alternative to ChIP-seq for mapping genome-wide BMAL1 occupancy.

**Figure 1:**
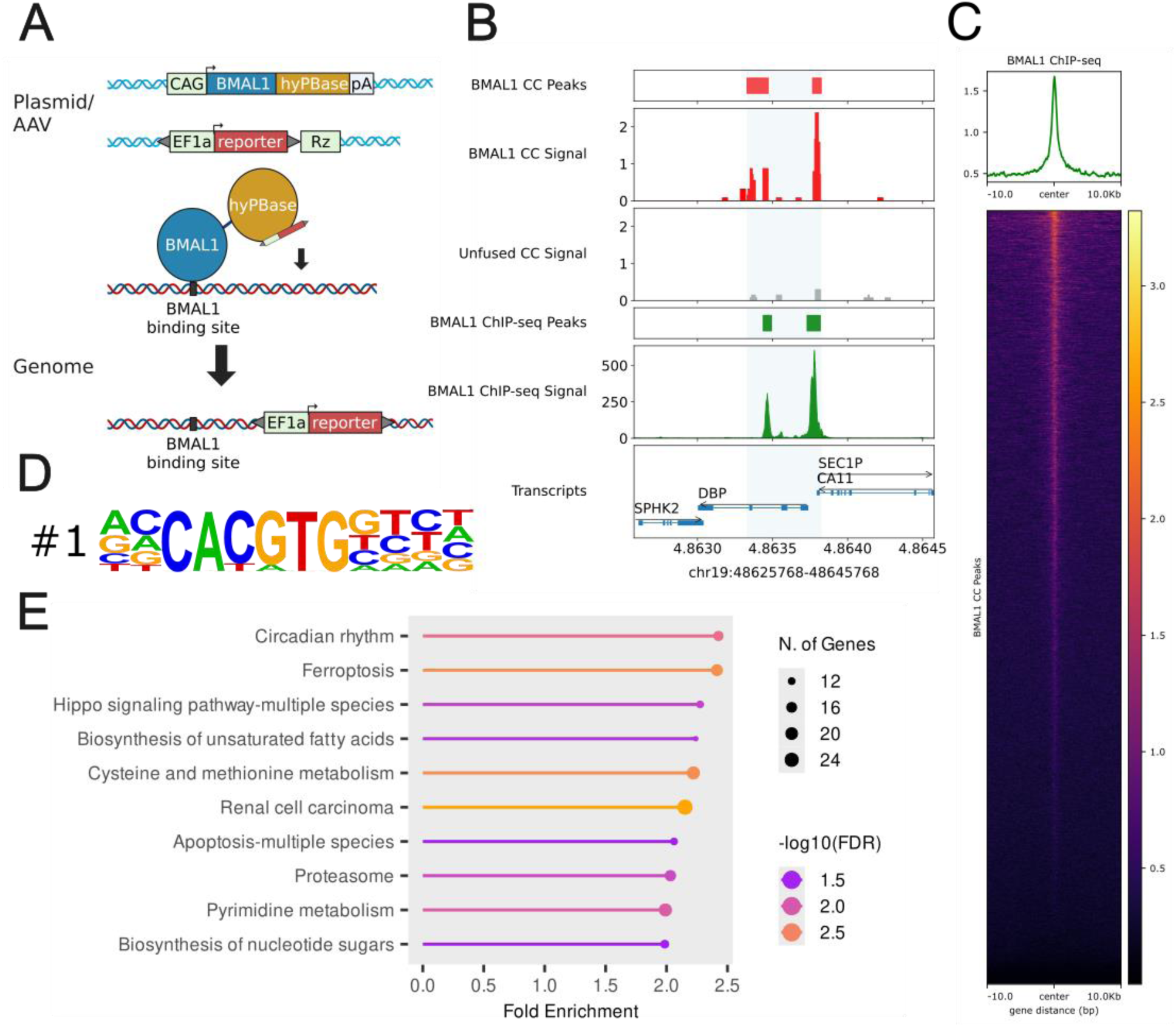
The Calling Cards (CC) technology accurately records BMAL1 binding in HepG2 cells. A) Schematic of the CC assay: a CAG-driven BMAL1-hyPBase fusion and an EF1α-driven self-reporting transposon reporter are delivered by either plasmid or AAV; reporter insertions deposited near BMAL1 binding sites are recovered by performing the CC library prep protocol and then sequencing. (B) Genome browser example showing BMAL1 CC peaks and insertion signal (red), unfused hyPBase control signal (gray), and BMAL1 ChIP-seq peaks and signal (green), illustrating concordant enrichment. (C) BMAL1 ChIP-seq profile plot and heatmap centered on BMAL1 CC peaks (±10 kb from center), demonstrating ChIP-seq enrichment at the CC peak centers. (D) Motif enrichment from HOMER analysis of BMAL1 CC peaks identifies the canonical E-box (CACGTG), consistent with BMAL1 binding. (E) ShinyGO KEGG enrichment for genes associated with BMAL1 CC peaks (nearest gene / within 5 kb); dot size indicates the number of genes per term and color indicates -log10(FDR), with circadian rhythm as the top enriched pathway.

### Combining Calling Cards with MACS enables astrocyte-specific BMAL1 binding profiles

We next asked whether Calling Cards could be combined with astrocyte purification from brain homogenates to obtain astrocyte-enriched BMAL1 binding profiles. We used magnetic-activated cell sorting (MACS), in which dissociated brain cells are incubated with an antibody against a cell-surface antigen conjugated to magnetic beads; labeled cells are retained on a magnetic column while unlabeled cells flow through, and the bound fraction is eluted after removal from the magnet. Our MACS protocol, adapted from published approaches (27), sequentially removes myelin debris and dead cells and then enriches for astrocytes using ACSA2 selection; remaining oligodendrocytes and neurons were depleted in the flow-through. Our protocol for MACS-CC then combines MACS sorting with Calling Cards, in which mice are injected with CC AAVs at age P0-P1, then the Calling Card constructs record BMAL1 binding over 2 months, when the mice are euthanized and the brains are sorted for astrocytes using MACS. We then extract RNA, perform CC library prep, and sequence (Figure 2A).

**Figure 2:**
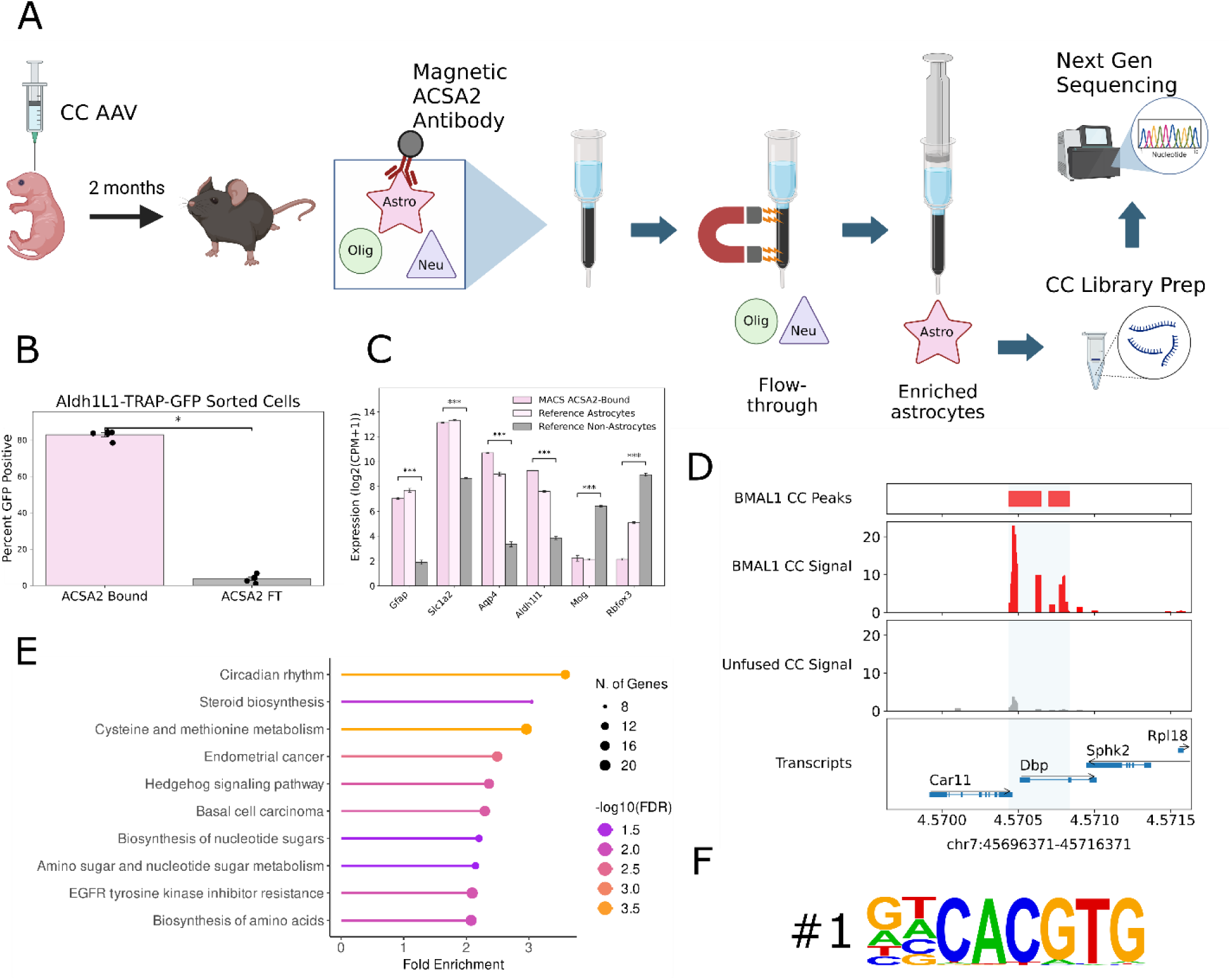
MACS-Calling Cards maps BMAL1 occupancy specifically in astrocytes. (A) Workflow: mice are injected with Calling Cards (CC) AAV, and after 2 months brains are dissociated and astrocytes are enriched by ACSA2 magnetic-activated cell sorting (MACS) prior to CC library preparation and next-generation sequencing. (B) MACS enriches astrocytes: percentage of Aldh1L1-TRAP-GFP+ cells in the ACSA2-bound and flow-through (FT) fractions (points =individual mice) Significance test = Exact two-sided permutation test on the difference in means. Adjusted Pval *Gfap* = 7.24E-06, adjusted Pval *Slc1a2* = 1.11E-12, adjusted Pval *Aqp4* = 1.30E-05, adjusted Pval *Aldh1l1* = 2.13E-05, adjusted Pval *Mog* = 2.67E-11, adjusted Pval *Rbfox3* = 9.30E-09. (C) Marker expression confirms astrocyte enrichment in the ACSA2-bound fraction (astrocyte markers Gfap, Slc1a2, Aqp4, Aldh1l1) with depletion of non-astrocyte markers (Mog, oligodendrocyte; Rbfox3/NeuN, neuronal), compared to reference astrocyte and non-astrocyte expression from a published snRNA-seq atlas. Significance test = Welch’s two-sample t-test (D) Genome browser example (chr7:45,701,371–45,711,371) showing MACS2-called BMAL1 CC peaks and CC insertion signal in ACSA2-bound cells (red) versus low background signal from the unfused hyPBase control (gray). (E) ShinyGO KEGG enrichment for genes associated with BMAL1 CC peaks (nearest gene / within 5 kb); dot size indicates number of genes per term and color indicates -log10(FDR). (F) HOMER motif enrichment of BMAL1 CC peaks identifies the canonical E-box (CACGTG) as the top enriched motif.

We validated our astrocyte sorting method with MACS by using Aldh1l1-Rpl-GFP TRAP mice, in which GFP specifically marks astrocytes. Flow cytometry showed that the ACSA2-bound fraction was strongly enriched for GFP+ cells (∼80%), whereas the flow-through contained <10% GFP+ cells (Figure 2B). RNA-seq of the bound fraction confirmed upregulation of astrocyte markers (including *Gfap, Aqp4*, and *Aldh1l1*) and depletion of neuronal (*Rbfox3*) and oligodendrocyte (*Mog*) markers relative to a single-nucleus reference atlas (Figure 2C) (35).These results indicate that from brain homogenates the MACS protocol yields astrocyte-enriched populations suitable for downstream assays

We then combined MACS with BMAL1 Calling Cards (MACS-CC) to measure BMAL1 binding in astrocytes *in vivo*. In MACS-isolated astrocytes, MACS-CC identified BMAL1 peaks at canonical circadian targets such as *Dbp* (Figure 2D). Genes near MACS-CC peaks were enriched for circadian rhythm categories (Figure 2E) and motif analysis recovered the E-box motif (Figure 2F). Thus, Calling Cards can be integrated with astrocyte enrichment to enable cell-type-enriched mapping of BMAL1 occupancy in the brain.

### BMAL1 binding is reprogrammed in *Cln3^Δex7/8^* astrocytes

Having validated our MACS-CC technique, we tested BMAL1 binding in astrocytes isolated from 2-month-old WT or *Cln3^Δex7/8^* mice. We chose this age because it represents an early state of pathology that is still targetable by AAV and allows for high throughput experiments (21, 22, 36). We mapped BMAL1 occupancy genome-wide and identified sites that were WT-specific, *Cln3^Δex7/8^*-specific, or shared (Figure 3A). Peak calling and gene assignment were performed with PyCallingCards to define shared, WT-specific, and *Cln3^Δex7/8^*-specific target gene sets. BMAL1 binding was extensively redistributed, with comparable numbers of WT-specific and *Cln3^Δex7/8^*-specific peaks (with 1500 CLN3-specific peaks, 1498 WT-specific peaks, and 16387 Shared peaks) these were nearby 820 unique genes for CLN3-specific, 862 unique genes for WT-specific, and 9406 genes for shared (Figure 3B, visualized genome-wide by Figure 3C). To evaluate functional relevance, we leveraged astrocyte-specific circadian expression data (8) and compared rhythmicity metrics for genes near each peak class. Genes closest to WT-specific and *Cln3^Δex7/8^*-specific BMAL1 peaks had significantly lower FDR values for circadian rhythmicity than genes near shared peaks, with WT-specific genes having the lowest adjusted p values (Figure 3D). This suggests BMAL1’s remodeling happens preferentially at loci where transcription is robustly rhythmic. We plotted the significantly circadian genes for each category (ranked by phase) and noticed that genes nearby WT-specific peaks had noticeably more extreme troughs (such as those around 8 hours) and peaks (such as those around 22 hours) (Figure 3E). In order to determine if condition-specific genes exhibit higher amplitudes than shared peaks, we calculated the average z-scores for expression for hours near the peak (-2 hours from peak, peak hour, and +2 hours from peak), and for hours mid cycle (+10, +12, and +14 hours from peak) for genes of each category that are significantly circadian. We observed that WT-specific peaks demonstrated a more extreme difference between (Figure 3F). These results indicate that genes closest to WT-specific peaks are more circadian (lower adjp) and have more extreme oscillations than shared peaks.

**Figure 3:**
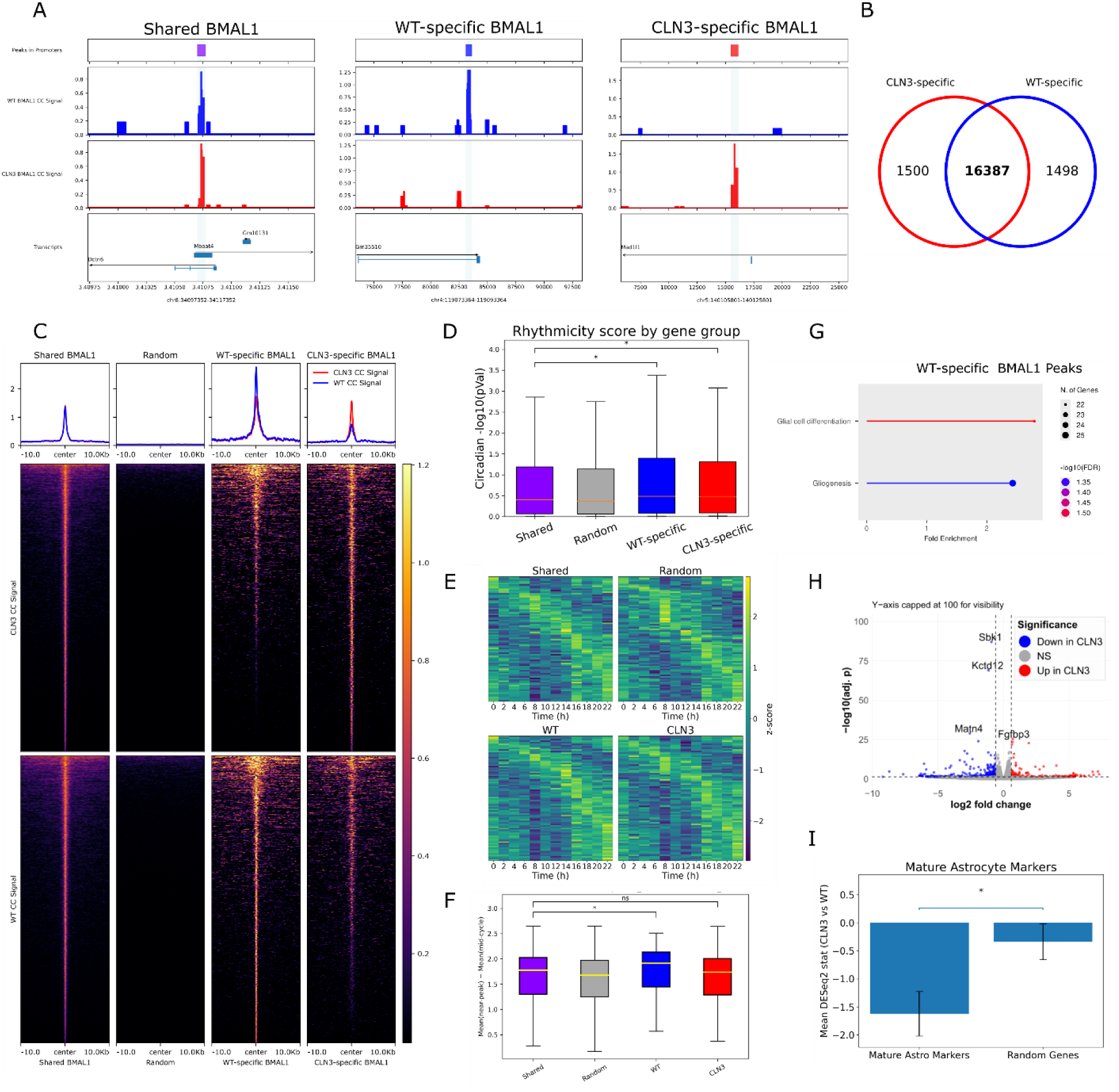
BMAL1 occupancy is extensively redistributed in *Cln3^Δex7/8^* astrocytes. Using MACS-enriched astrocytes and BMAL1 Calling Cards, we mapped genome-wide BMAL1 occupancy in WT (blue) and *Cln3^Δex7/8^* (red) astrocytes and classified peaks as shared, WT-specific, or *Cln3^Δex7/8^*-specific. (A) Genome browser examples of each peak class showing WT and *Cln3^Δex7/8^*BMAL1 CC signal and the corresponding peak interval (shaded). (B) Peak overlap summary and peak-to-gene assignment (PyCallingCards) defining shared (16,387), WT-specific (1,498), and *Cln3^Δex7/8^*-specific (1,500) BMAL1 peaks.(C) Aggregate profiles and heatmaps of WT and *Cln3^Δex7/8^*BMAL1 CC signal centered on shared peaks, WT-specific peaks, *Cln3^Δex7/8^*-specific peaks, and matched random regions (±10 kb). (D) Circadian rhythmicity of genes nearest each peak class using an astrocyte circadian expression reference dataset (Sheehan et al., 2025), plotted as rhythmicity score (-log10 adjusted rhythmicity metric); genes associated with WT-specific and *Cln3^Δex7/8^*-specific peaks show significantly stronger rhythmicity than genes associated with shared peaks. Significance test = Mann-Whitney U comparing Shared vs WT-specific or *Cln3^Δex7/8^*-specific. Shared vs. WT-specific pvalue = 1.505e-02; Shared vs. *Cln3^Δex7/8^*-specific pvalue = 1.505e-02. (E) Phase-normalized heatmap of top 150 average expression levels (z-scored) at near-peak hours (-2,0, and +2 hours around peak) and mid-cycle hours (+10,+12,+14 hours from peak). Genes first downsampled to the same sample size for all and then ranked by difference in mean z-score between near-peak and mid-cycle. (F) Boxplot summary of genes represented in E; Shared vs. WT pvalue = 0.026; Shared vs *Cln3^Δex7/8^*-specific pvalue = 0.63; significance test = Mann-Whitney U test. (G) ShinyGO’s GO Biological Process enrichment for genes near WT-specific BMAL1 peaks highlights glial differentiation-related terms, whereas genes near *Cln3^Δex7/8^*-specific peaks do not yield coherent enrichment, consistent with diffuse retargeting. Background for each is genes nearby shared peaks. (H) Differential gene expression from RNA-seq of MACS astrocytes (*Cln3^Δex7/8^* vs WT; |log2FC| > 0.6, adj. p < 0.05). (I) Mature astrocyte marker genes are significantly reduced in *Cln3^Δex7/8^* astrocytes compared with a matched random gene set. Significance test = Two-sided permutation test. P = 0.001.

We next sought to ask whether WT-specific and *Cln3^Δex7/8^*-specific peaks differed in the functional categories of genes they were bound to. WT-specific BMAL1 peaks were enriched near genes related to glial differentiation, whereas *Cln3^Δex7/8^*-specific peaks yielded no significant enrichment categories (Figure 3G), consistent with a more diffuse retargeting model. These results are consistent with BMAL1 losing coherent functional targeting in disease. To test whether the loss of WT-like BMAL1 targeting is reflected in the astrocyte transcriptome, we performed RNA-seq on WT and *Cln3^Δex7/8^*astrocytes. BMAL1 has been implicated in neural development, and BMAL1 loss can alter astrocytic differentiation outcomes *in vitro* (37–39), raising the possibility that disrupted BMAL1 binding contributes to altered maturation. We identified 341 differentially expressed genes (|log2FC| > 0.6, adjp < 0.05; Figure 3H). Cytokine signaling was the top enriched category (Supplementary Figure S1A), and many cytokine-associated transcripts were downregulated in *Cln3^Δex7/8^* astrocytes (Supplementary Figure S1B), consistent with decreased cytokine/chemokine secretion reported in this model (21). Notably, mature astrocyte marker genes were significantly reduced in *Cln3^Δex7/8^*astrocytes (Figure 3I), linking BMAL1 binding changes to a less mature transcriptional profile.

Together, these data indicate that WT-specific BMAL1 binding events are preferentially associated with circadian and glial differentiation programs, and that *Cln3^Δex7/8^* astrocytes display reduced expression of maturity markers. In contrast, newly acquired *Cln3^Δex7/8^*-specific BMAL1 peaks do not cluster into any obvious functional gene category.

### Chromatin accessibility is reprogrammed in *Cln3^Δex7/8^* astrocytes

Given the extensive redistribution of BMAL1 binding and its transcriptional associations in *Cln3^Δex7/8^* astrocytes, we asked what mechanisms drive BMAL1 reprogramming. In other papers, BMAL1 occupancy is aligned with tissue-specific open chromatin (40), suggesting that chromatin accessibility restricts BMAL1 binding. Therefore, we wanted to test whether these findings apply to BMAL1 binding in CLN3 disease. To determine the extent to which altered chromatin accessibility contributes to BMAL1 retargeting in *Cln3^Δex7/8^* astrocytes, we profiled promoter architecture in the MACS-enriched astrocytes using MNase-seq. MNase-seq maps nucleosome positions by using micrococcal nuclease to preferentially cut exposed “linker” DNA between nucleosomes, then sequencing the protected DNA fragments to infer where nucleosomes sit across the genome. We performed MNase-seq in astrocytes sorted using our MACS protocol. To assess data quality, we visualized MNase-seq signal around annotated transcription start sites (TSS) and observed the expected TSS-centered depletion with phased flanking nucleosomes in both genotypes, whereas random genomic positions showed no such structure (Supplementary Figure S2). We next formally identified disease-specific NFRs by defining, for each annotated TSS, a strand-aware 150-bp putative NFR window flanked by two 150-bp windows. An NFR was called when the mean flanking signal was ≥0.5 CPM and at least 1.3-fold higher than the signal within the putative NFR window, consistent with well-positioned −1/+1 nucleosomes flanking a depleted TSS.

Using this framework, we observed extensive remodeling of promoter accessibility between WT and *Cln3^Δex7/8^*astrocytes. Genome-browser examples illustrate each class (Figure 4A): shared NFRs show low MNase-seq signal at the TSS in both genotypes, WT-specific NFRs exhibit a clear TSS depletion in WT that is lost in disease, and *Cln3^Δex7/8^*-specific NFRs show a reciprocal pattern - depletion that is absent in WT. When aggregated across all promoters, average MNase-seq profiles and heatmaps around TSSs recapitulated these patterns (Figure 4B). Promoters classified as “Both NFR” displayed a pronounced trough at the TSS in both WT and *Cln3^Δex7/8^* astrocytes, whereas promoters classified as “Neither NFR” retained high signal at the TSS in both conditions. In contrast, condition-specific categories showed genotype-restricted TSS depletion: WT-specific NFRs had a trough selectively in WT, while *Cln3^Δex7/8^*-specific NFRs had a trough selectively in *Cln3^Δex7/8^* astrocytes (Figure 4B). Consistent with this widespread remodeling, we identified thousands of promoter NFRs that were shared (8100) as well as large condition-specific sets (2402 WT-specific and 2343 CLN3-specific; Figure 4C). Figure 4D summarizes the distribution of accessibility signals across those peak categories (Figure 4D). Computing the between-genotype difference in MNase signal showed that shared NFRs (and matched random controls) were centered near zero, whereas WT-specific and CLN3-specific NFRs shifted in opposite directions and differed significantly from shared promoters (Figure 4E). Together, these results indicate that promoter-proximal chromatin architecture is broadly altered in *Cln3^Δex7/8^* astrocytes.

**Figure 4:**
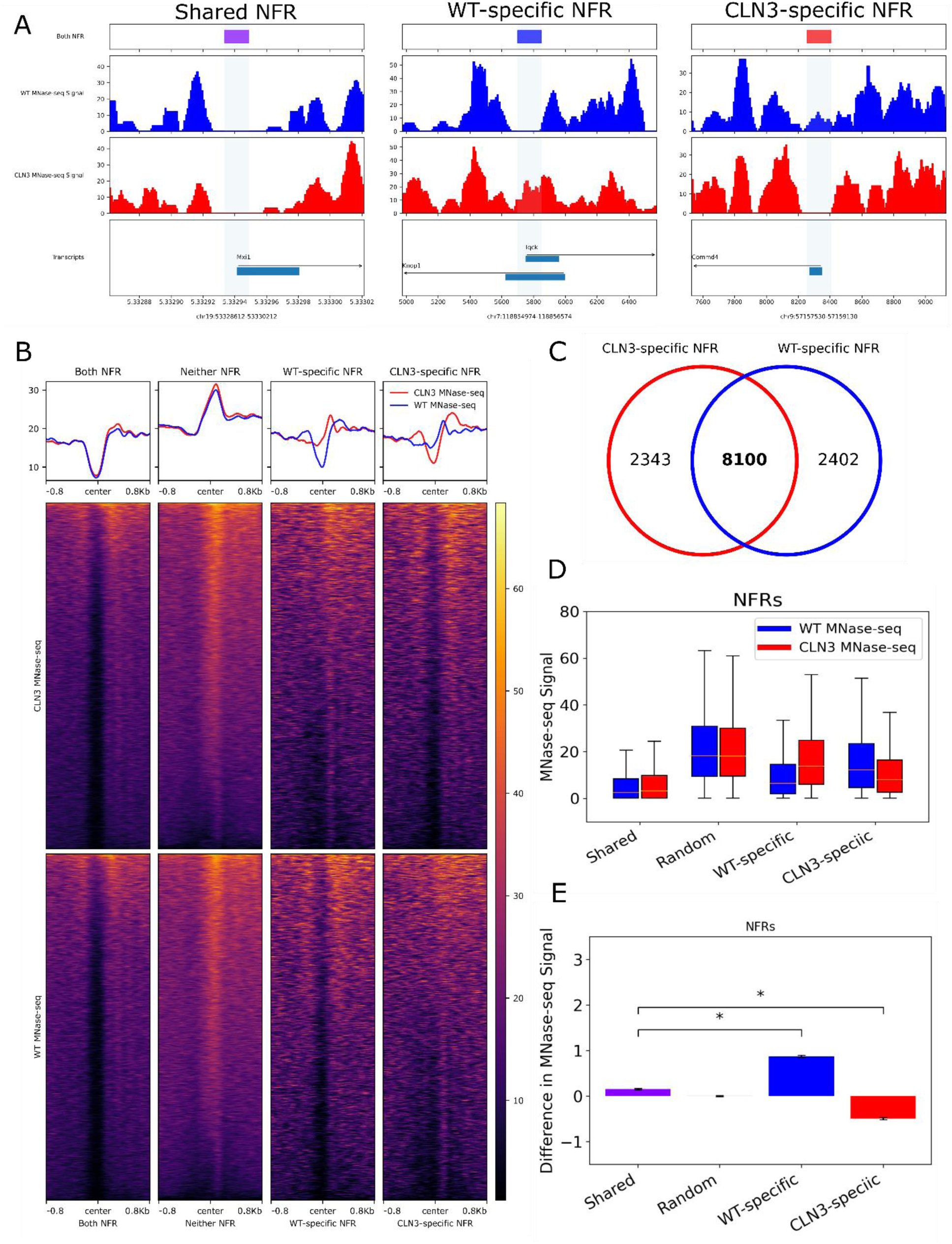
Promoter-proximal nucleosome-free regions are extensively remodeled in *Cln3^Δex7/8^* astrocytes. (A) Genome-browser examples of a shared NFR, a WT-specific NFR, and a CLN3-specific NFR, showing MNase-seq signal in WT (blue) and *Cln3^Δex7/8^*(red) at promoter regions; the shaded region marks the TSS-centered NFR window and gene models are shown below. (B) Aggregate MNase-seq profiles (top) and corresponding heatmaps (bottom) across promoters stratified into “Both NFR,” “Neither NFR,” “WT-specific NFR,” and “CLN3-specific NFR” categories, plotted ±0.8 kb around the TSS center for WT and *Cln3^Δex7/8^*astrocytes. (C) Venn diagram showing overlap of promoter NFR calls between genotypes (counts indicated). (D) Distribution of MNase-seq signal within the promoter NFR window across categories for WT (blue) and *Cln3^Δex7/8^*(red); “Random” denotes matched control windows. (E) Mean genotype difference in MNase-seq signal for each category (ΔMNase = *Cln3^Δex7/8^*− WT), with brackets indicating the statistical comparisons shown (*, significant; ns, not significant). Significance test = two-sided Mann-Whitney U test. Adjusted pvalue shared vs. WT-specific = 2.40E-163, adjusted pvalue shared vs. CLN3-specific = 9.84E-128. Promoter NFRs were defined using a strand-aware 150-bp TSS-centered window flanked by two 150-bp windows; an NFR was called present when mean flanking signal was ≥0.5 CPM and ≥1.3-fold higher than signal within the NFR window.

### BMAL1 reprogramming is uncoupled from chromatin accessibility in *Cln3^Δex7/8^* astrocytes

To test whether such altered chromatin accessibility explains BMAL1 retargeting within CLN3 deficient astrocytes, we integrated BMAL1 peak calls from WT and *Cln3^Δex7/8^* astrocytes with matched MNase-seq measurements of chromatin openness. We first utilized our BMAL1 peaks that have been categorized as shared peaks, WT-specific peaks, and *Cln3^Δex7/8^*-specific peaks and asked a straightforward question: at BMAL1 sites that are gained or lost between genotypes, is chromatin accessibility also gained or lost in the same direction? In Figure 5A, average MNase-seq signal is plotted across ±5 kb around each BMAL1 peak set, with WT and *Cln3^Δex7/8^* MNase profiles overlaid and a matched random set shown as a negative control. Shared BMAL1 peaks show the expected local accessibility signature (a pronounced trough centered on the peak), and importantly, WT-specific and *Cln3^Δex7/8^*-specific BMAL1 peaks also sit in comparably open chromatin in both genotypes - the MNase profiles for WT and CLN3 are largely superimposable even at peaks that are classified as condition-specific (i.e. the blue and red lines overlap). Figure 5B summarizes this same comparison by quantifying MNase-seq signal distributions across the peak categories, again showing minimal genotype-dependent shifts in accessibility at shared and condition-specific BMAL1 peaks relative to the random control. Finally, to move beyond discrete peak classes, we computed per-peak ΔAccessibility and ΔBMAL1 binding across the union of all BMAL1 peaks (shared + WT-specific + *Cln3^Δex7/8^*-specific) and tested whether changes in accessibility predict changes in BMAL1 occupancy genome-wide. This analysis (Figure 5C) shows essentially no relationship between ΔAccessibility and ΔBMAL1 (Spearman ρ ∼ 0; slope ∼ 0), indicating that accessibility differences do not systematically explain where BMAL1 is gained or lost in *Cln3^Δex7/8^* astrocytes. Notably, the absence of correlation is driven by the same feature emphasized in panels A-B: there is little to no consistent change in accessibility between WT and *Cln3^Δex7/8^* at BMAL1 peak regions, including at condition-specific BMAL1 peaks, yet BMAL1 occupancy is still redistributed, supporting a model in which within-cell type BMAL1 reprogramming can occur without coordinated chromatin opening/closing, instead perhaps relying on other factors, such as context-specific interactions with co-binders.

**Figure 5:**
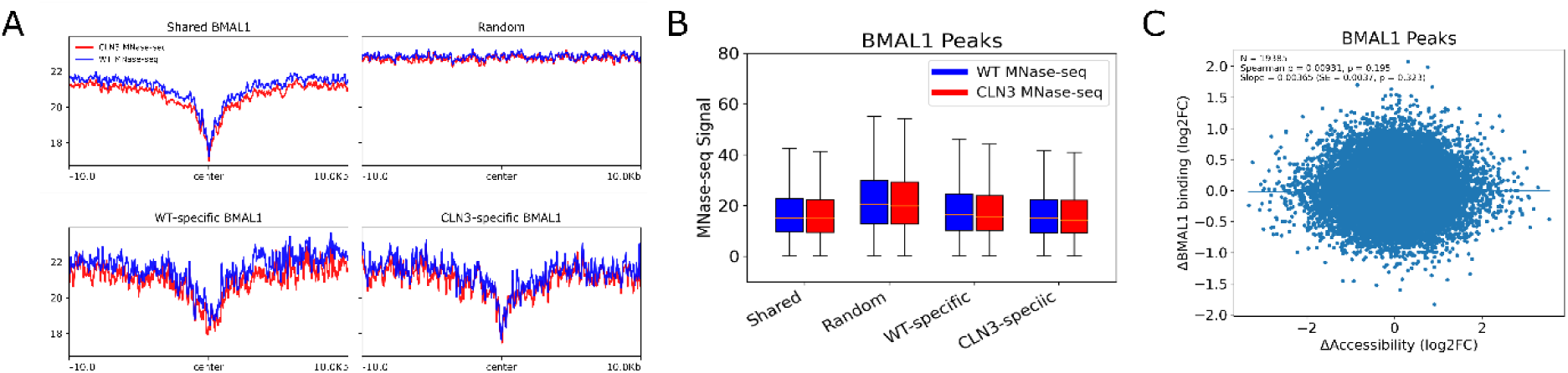
Chromatin accessibility is similar at shared, WT-specific, and *Cln3^Δex7/8^*–specific BMAL1 binding sites. (A) Average MNase-seq profiles (±10 kb from peak center) in WT (blue) and *Cln3^Δex7/8^* (red) astrocytes at shared, WT-specific, and CLN3-specific BMAL1 peaks, plus random matched control regions (“Random”). (B) Boxplots of MNase-seq signal summarized in a 500 bp window around peak centers show comparable MNase-seq signal between genotypes across all peak classes. (C) Across all BMAL1 peaks (N = 19,385), genotype-dependent changes in MNase-seq signal (ΔAccessibility = log2(WT/*Cln3^Δex7/8^*)) do not predict changes in BMAL1 occupancy measured by calling cards (ΔBMAL1 binding = log2(WT/*Cln3^Δex7/8^*), indicating little-to-no coupling between accessibility changes and BMAL1 binding changes in this dataset (Spearman ρ and linear fit shown). Significance test = Spearman rank correlation and linear regression slope test.

### Chromatin accessibility does not consistently explain BMAL1 reprogramming in other settings

Across tissues, BMAL1 binding has been reported to track with the accessible chromatin landscape: tissue-specific BMAL1 peaks tend to occur at tissue-specific open regions, suggesting that differences in accessibility can be a major driver of apparent “reprogramming” (40). In contrast, in *Cln3^Δex7/8^* astrocytes we found that differential accessibility does not explain where BMAL1 is gained or lost, indicating that within-cell type perturbations may decouple BMAL1 occupancy from condition-dependent changes in accessibility. To place this result in the context of prior BMAL1 reprogramming studies, we reanalyzed published datasets in which BMAL1 binding changes could be directly compared to matched accessibility measurements. Specifically, we analyzed: 1) BMAL1 ChIP-seq comparing liver and kidney (GSE110604, (40)) with ATAC-seq data from the same tissues (PRJNA497970, (41); 2) BMAL1 ChIP-seq comparing normal chow (NC) and high-fat diet (HFD) (GSE117488, (42)) with FAIRE-seq under NC and HFD (GSE75984, (43)); and 3) BMAL1 ChIP-seq comparing ad libitum (AL) and calorie restriction (CR) feeding (GSE151281, (44)) with ATAC-seq comparing AL and CR (CRA010174, (45). For each comparison, we first called BMAL1 peaks in each condition and partitioned them into shared peaks (present in both) and condition-specific peaks (present only in one condition), and then asked a focused question: at BMAL1 peaks that are condition-specific, is chromatin accessibility also condition-specific? We plotted the average ATAC-seq/FAIRE-seq signal centered on BMAL1 peak summits for shared peaks, a random control set, and each condition-specific peak set, overlaying the two conditions’ accessibility tracks (Figure 6A, with the quantities of condition-specific and shared peaks in Figure 6B). We then summarized the distribution of accessibility signals across those peak categories (Figure 6C). Liver-versus-kidney shows pronounced, directionally matched accessibility differences at tissue-specific BMAL1 peaks (e.g., liver-specific BMAL1 peaks sit in more accessible chromatin in liver than kidney, and vice versa), whereas in the HFD-versus-NC comparison, the FAIRE-seq profiles are nearly superimposable even at HFD-specific and NC-specific BMAL1 peaks, indicating that BMAL1 gains/losses occur with essentially no accompanying change in chromatin accessibility. Finally, we quantified coupling genome-wide by computing per-peak ΔAccessibility and ΔBMAL1 binding across the union of all BMAL1 peaks (shared + condition-specific) and correlating these values (Figure 6D). Consistent with the signal plots, we observe strong coupling for liver-versus-kidney, no coupling for HFD-versus-NC, and a modest relationship for CR-versus-AL. Specifically, Spearman correlations across the union peak set were ρ=0.759 (p=3.99×10⁻³¹²; N=1661) for liver vs. kidney, ρ=0.00764 (p=0.517; N=7189) for HFD vs. NC, and ρ=0.286 (p<1×10⁻³⁰⁰; N=19,943) for CR vs. AL. Importantly, the lack of correlation in HFD-versus-NC is expected precisely because ΔAccessibility has little dynamic range in this dataset (accessibility is largely unchanged between diets, including at diet-specific BMAL1 peaks). This mirrors our *Cln3^Δex7/8^* astrocyte data: BMAL1 occupancy can be reprogrammed in the absence of corresponding accessibility remodeling, supporting a model in which within-tissue BMAL1 retargeting can reflect mechanisms other than chromatin opening/closing (e.g., altered co-factor engagement). Together, these analyses suggest that chromatin accessibility is a dominant determinant of BMAL1’s tissue-specific cistrome - where different tissues present genuinely different baseline open-chromatin landscapes - but it is not a reliable predictor of BMAL1 retargeting across external perturbations within a tissue or cell type, where accessibility can remain largely stable even as BMAL1 occupancy shifts. In other words, tissue-to-tissue comparisons may look “accessibility-driven” because the substrate differs, whereas condition-driven reprogramming can occur through mechanisms that change BMAL1 recruitment without requiring broad remodeling of chromatin openness.

**Figure 6:**
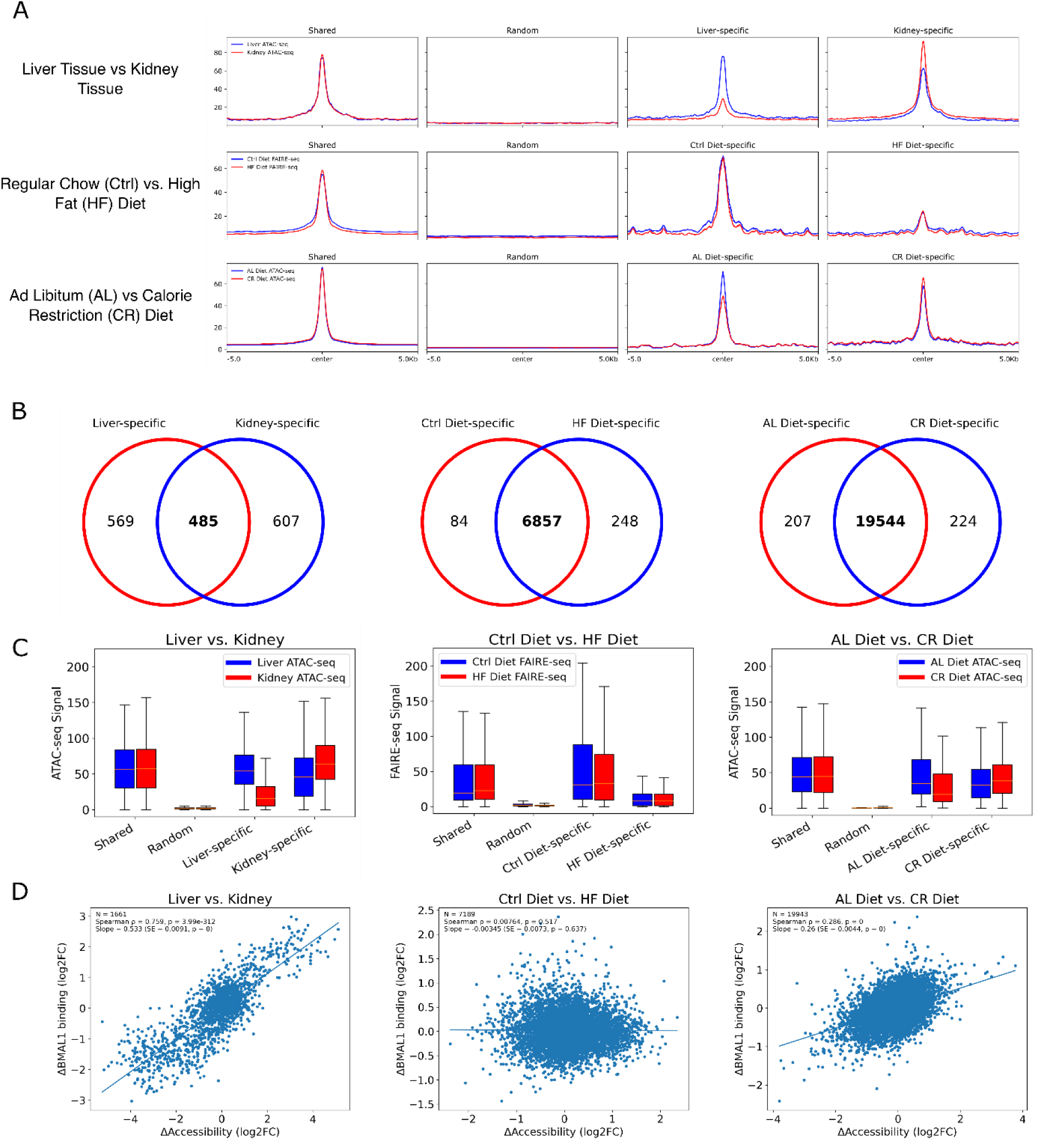
Published BMAL1 reprogramming datasets show context-dependent coupling between chromatin accessibility and BMAL1 occupancy. We re-analyzed three prior BMAL1 perturbation/reprogramming paradigms with matched BMAL1 ChIP-seq and accessibility profiling (ATAC-seq or FAIRE-seq): liver vs kidney, control/normal chow vs high-fat diet (HFD), and ad libitum (AL) vs calorie restriction (CR). (A) Aggregate accessibility signal (ATAC-seq or FAIRE-seq) centered on BMAL1 ChIP-seq peak centers (±3 kb), shown for shared peaks, condition-specific peaks, and matched random control regions. (B) Overlap of BMAL1 peak sets between conditions for each comparison. (C) Distribution of mean accessibility signal within a 500 bp window around BMAL1 peak centers (±250 bp) for each peak class, plotted separately for each condition. (D) For each BMAL1 peak, change in accessibility (ΔAccessibility, log2FC of ATAC/FAIRE signal; kidney/liver, HFD/control, or CR/AL) plotted against change in BMAL1 binding (ΔBMAL1 binding, log2FC of BMAL1 ChIP–seq signal for the same comparison). Significance test = Spearman rank correlation and linear regression slope test. Each point represents one peak; the fitted line and Spearman correlation (ρ) are shown in-panel.

### Disrupted cofactor-associated motif architecture accompanies BMAL1 retargeting in *Cln3^Δex7/8^* astrocytes

Because chromatin accessibility did not correlate with reprogramming of BMAL1 binding, we wondered if differential cofactors association better explained BMAL1 reprogramming. We quantified cofactor association through motif occupancy. We hypothesized that if a cofactor was more active in either WT or *Cln3^Δex7/8^* astrocytes, we would see enrichment of that motif in the corresponding condition-specific NFRs. Accordingly, if that cofactor associated with BMAL1 in a condition specific manner, we should see an enrichment of that motif nearby BMAL1 motifs in a similar condition-specific manner. We therefore analyzed (i) condition-specific open promoters (chromatin-defined) and (ii) condition-specific BMAL1 peaks at promoters (binding-defined) and tested each set for enriched motif occupancy (Figure 7A-B). Condition specific promoters were defined based on extending the boundaries of our previous condition-specific NFRs to 2kb around the TSS. We considered motifs as “passing” if they were significant in both analyses. In WT astrocytes, WT-specific open promoters and WT-specific BMAL1 promoter targets showed numerous significantly enriched motifs with positive effect sizes (FDR < 0.05) (Figure 7A), including STAT- and TEAD-like signatures. In contrast, there were no motifs significant in both *Cln3^Δex7/8^*-specific open promoters and *Cln3^Δex7/8^*-specific BMAL1 peaks at promoters (Figure 7B), suggesting a collapse of coherent motif structure in CLN3 disease-defined promoter sets. The top five WT-enriched motifs from each WT-specific analysis are shown as sequence logos (Figure 7C). We then aggregated candidate TFs predicted to bind these top WT motifs (across both WT-specific promoter sets) and examined their expression changes. We took a generous approach, noting how motifs such as PRDM14 potentially contain the AGGTCT nuclear receptor (NR) half-site motif variant and how the TEAD2 motif also contains the GGAA ETS core (Supplementary Table S6 for full gene list). None of the candidate TFs were expressed to be counted as a DEG (FDR<0.05, log2fc > 0.6), but Rxra, Stat3, and Nr1d2 had FDR values <0.05 (Figure 7D). These three TFs are therefore the most likely cofactors of BMAL1 to explain WT-specific binding.

**Figure 7:**
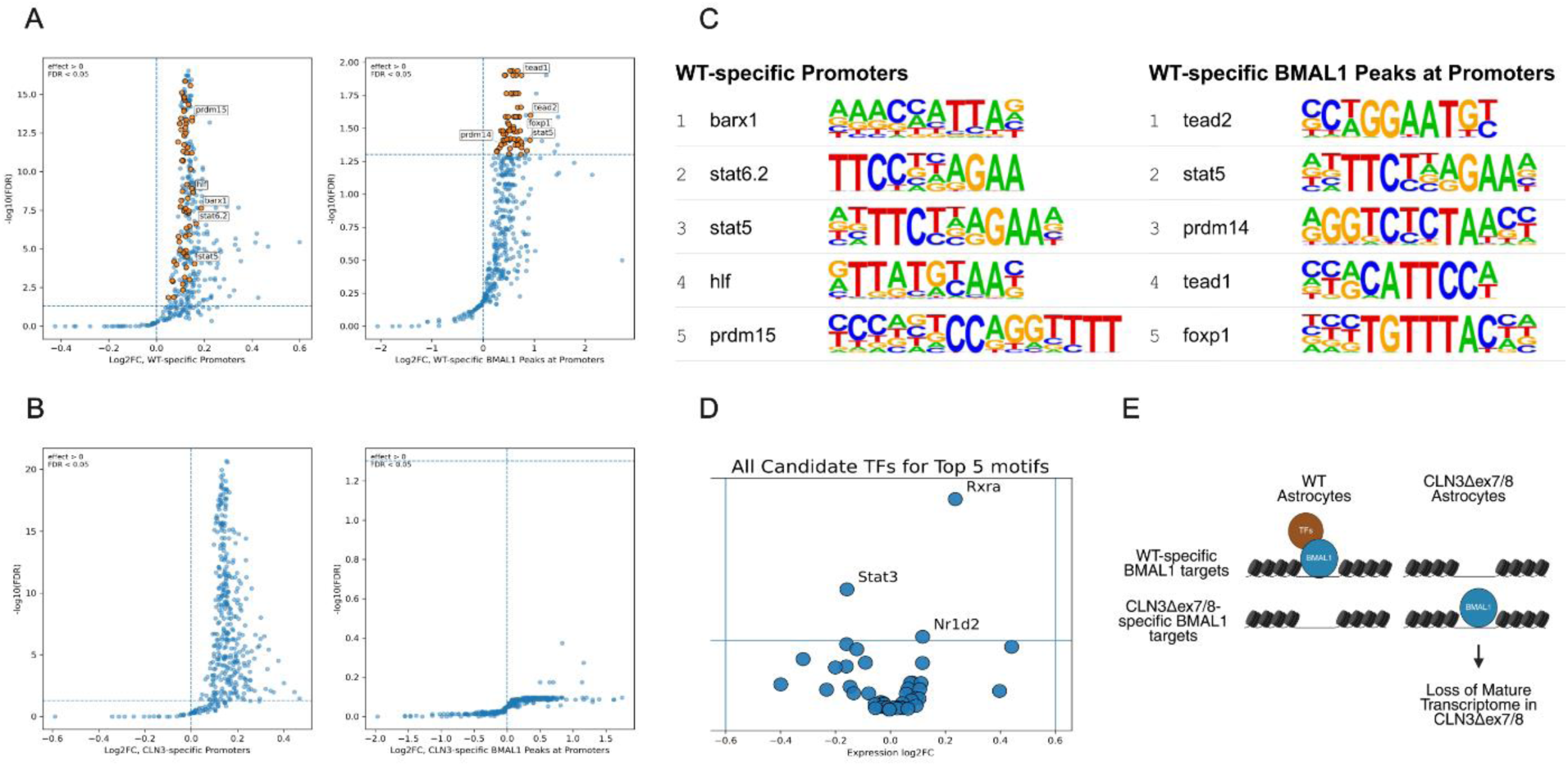
Disrupted cofactor-associated motif architecture accompanies BMAL1 retargeting in *Cln3^Δex7/8^*astrocytes. (A-B) Differential motif-occupancy enrichment analyses for (i) condition-specific open promoters (chromatin-defined; promoters defined by extending condition-specific NFR boundaries to ±2 kb around the TSS) and (ii) condition-specific BMAL1 peaks at promoters (binding-defined). For each motif, the effect size (log2FC enrichment) is plotted against significance (-log10 FDR); dashed lines mark effect = 0 and FDR = 0.05, and labeled points highlight representative enriched motifs. WT-specific promoter sets show multiple significantly enriched motifs with positive effect sizes (FDR < 0.05), including STAT5/STAT6-, and TEAD-like motifs (A). In contrast, *Cln3^Δex7/8^*-specific promoter sets fail to show motifs that are significant in both the open-promoter and BMAL1-promoter-peak analyses (B), indicating loss of coherent motif structure in disease-defined promoter targets. (C) Sequence logos for the top five WT-enriched motifs from each WT-specific analysis (open promoters, left; BMAL1 peaks at promoters, right). (D) Expression changes (log2FC, *Cln3^Δex7/8^*vs WT) for candidate TFs predicted to bind the top WT-enriched motifs (aggregated across both WT-specific promoter analyses), plotted against -log10(padj); Rxra, Stat3, and Nr1d2 are highlighted as significant by padj (< 0.05) but do not meet DEG thresholds used here (padj < 0.05 and |log2FC| > 0.6). (E) Model: in WT astrocytes, cooperative cofactor networks help focus BMAL1 onto WT-specific circadian and glial differentiation gene programs; in *Cln3^Δex7/8^*astrocytes, weakening/loss of this motif-guided cofactor architecture leads to more diffuse BMAL1 targeting, reduced rhythmicity, and a shift away from mature astrocyte transcriptional states.

We propose a model in which BMAL1 retains the capacity to bind permissive/open chromatin in *Cln3^Δex7/8^* astrocytes, but chromatin accessibility alone does not specify where BMAL1 is targeted in CLN3 disease. Instead, loss or weakening of cooperative cofactor networks removes the WT-like motif-guided recruitment that normally focuses BMAL1 onto circadian and glial differentiation gene programs. As a result, BMAL1 binding becomes more diffuse and less programmatic in *Cln3^Δex7/8^* astrocytes, contributing to reduced rhythmicity and a shift away from mature astrocyte transcriptional states (Figure 7E).

## DISCUSSION

How the core clock transcriptional program is altered in neurodegenerative disorders remains a major question in circadian neurobiology. We focused on CLN3 disease and upon *Cln3^Δex7/8^*astrocytes because astrocyte dysfunction emerges early in disease progression in this neurodegenerative lysosomal storage disorder. We developed a new astrocyte-enriched *in vivo* BMAL1 occupancy assay (MACS-Calling Cards), and showed that BMAL1 binding is extensively reprogrammed in *Cln3^Δex7/8^* astrocytes. WT-specific BMAL1 sites preferentially map to circadian and glial differentiation programs, and *Cln3^Δex7/8^* astrocytes exhibit reduced expression of maturity markers. Despite widespread remodeling of promoter accessibility, differential accessibility does not explain gained or lost BMAL1 binding, supporting a model in which altered cooperative factor interactions, rather than accessibility, is a primary determinant of BMAL1 retargeting in disease.

A key advance of this study is MACS-CC, a tractable approach for mapping astrocyte-enriched TF occupancy in vivo. Calling Cards records binding while the animal is alive, reducing concerns that downstream purification steps perturb TF-DNA interactions - a limitation for ChIP-based methods that require cell harvesting prior to crosslinking. We used MACS-CC in this study to record BMAL1 binding, but MACS-CC can be used with any TF by fusing it to the hyPBase construct. Calling Cards has previously been used to characterize the binding of a variety of TFs (30). Other efforts to use Calling Cards in a cell type specific manner have involved using the Cre-lox system to flip a portion of hyPBase (31). This method adds additional basepair load to the AAV and, for larger TFs, can risk exceeding the limits of what can be packaged into AAV. The MACS-CC method works well with already validated Calling Card AAVs because the astrocyte sorting is a completely independent step.

Our study points to BMAL1 regulating astrocyte developmental genes within the context of a specific neurodegenerative disease, CLN3 disease or juvenile NCL. Evidence for the involvement of glia in the pathogenesis of multiple forms of NCL has accumulated recently (21, 36, 46–50). The early localized activation of either (or both) astrocytes and microglia corresponds closely to the distribution of subsequent neuron loss in the most vulnerable CNS regions (20, 51–55). This includes CLN3 disease, in which this glial activation appears morphologically compromised, taking a long time to reach a fully reactive state, but starting long before neurons die (21). As the NCL gene products are each expressed in both astrocytes and microglia, it is not surprising that glial dysfunction has been reported in multiple forms of NCL, including CLN3 disease. *In vitro* data from cultured CLN3-deficient astrocytes and microglia suggest their normal functions and responses to stimulation to be compromised, resulting in increased neuron loss in co-cultures (21). Most of the data on compromised glial function in CLN3 disease is centered in microglia, but our findings extend these observations to astrocytes.

Our findings from astrocytes isolated by MACS-CC reveal WT-specific BMAL1 peaks are enriched for glial differentiation genes, whereas *Cln3^Δex7/8^*astrocytes displayed a reduced maturity signature. This is consistent with the hypothesis that loss of WT-like BMAL1 targeting contributes to an altered astrocyte state in disease. This finding was somewhat surprising because BMAL1’s role in astrocytes is traditionally linked to astrocyte reactivity, as deletion of BMAL1 results in significantly increased GFAP expression (7, 14). This may be because BMAL1’s function in brain development is understudied, especially on a cell type-specific level, as previous reviews have noted (56, 57). However, several studies have found that BMAL1 does play a role in astrocyte development, as genetic disruption of *Bmal1* can bias neural stem/progenitor differentiation toward GFAP+ glial fates (38) and *Bmal1* deficiency early in life perturbs astrocyte structure and function *before* overt astrogliosis (58), suggesting that BMAL1 dysfunction can affect brain development before classical astrocyte activation becomes apparent. CLN3 disease is known to have altered brain development (59, 60), and several “adult” neurodegenerative diseases have been found to have a neurodevelopmental component (61–66). Our study provides further evidence that astrocytes in CLN3 disease display molecular changes early in life and suggests that altered BMAL1 binding contributes to these events long before neurodegeneration subsequently occurs.

We found that chromatin accessibility does not predict BMAL1 binding and found that, across other reprogramming contexts, there is not a consistent effect of chromatin accessibility on BMAL1 binding. This suggests accessibility can strongly shape BMAL1 occupancy in some settings, yet has inconsistent explanatory power for within-cell type perturbations, so the lack of correlation in *Cln3^Δex7/8^*astrocytes fits a broader pattern where partner networks and co-binding may matter more than baseline openness. This phenomenon has been observed with other TFs, such as MYC, where a study observed widespread changes in MYC occupancy without corresponding changes in chromatin accessibility at most MYC-bound sites, suggesting that TF occupancy can be extensively remodeled without broad accessibility changes (67).

Consistent with our chromatin accessibility finding, WT vs CLN3 disease motif analysis found >60 WT-enriched motifs and none enriched in disease. These results support a model in which loss of cooperative co-binder recruitment yields a more diffuse redistribution of BMAL1 across accessible sites without a dominant functional program. This interpretation fits well with the broader concept that BMAL1 binding is highly context dependent (varying by tissue and physiological state) and that co-binding TFs can be major determinants of where BMAL1 ultimately engages regulatory DNA, even when chromatin is accessible (40). Several of the enriched motif classes identified here (STAT-like, NR-like, and TEAD-like motifs) have precedent for influencing BMAL1 occupancy through shifts in partner-factor networks. First, STAT3, which is the dominant STAT family member associated with astrocyte differentiation and reactive astrocyte signaling (68, 69), also interacts with SMAD proteins in fetal neural progenitor cells in a cytokine stimulation-dependent manner, demonstrating that cellular context can reshape STAT-family interactomes and potentially alter if/how STAT proteins bind to BMAL1 in astrocytes (70, 71). Second, among NR-like factors, members of the core circadian NR REV-ERB family (which contains Nr1d2) are known to regulate clock output and shape circadian chromatin dynamics (72, 73), providing an established route by which changes in activity of those TFs could redirect BMAL1-regulated programs. In the liver, the NR HNF4A co-occupies many CLOCK:BMAL1 sites and can directly modulate CLOCK:BMAL1 transcriptional activity, linking NR activity to where BMAL1 binds (74). Third, for TEAD-like motifs, the TEAD co-activator YAP binds to BMAL1 in the epidermis as part of a BMAL1-YAP transcriptional complex, with TEAD likely forming part of this complex as well (75). Though none of the candidate TFs for our top 5 motifs passed the log2FC threshold to be counted as DEGs, post-transcriptional processing or altered binding to other cofactors could cause a co-binder of BMAL1 to reprogram BMAL1’s binding pattern.

Despite these caveats, our findings suggest a disease-context framework for circadian control in astrocytes. In *Cln3^Δex7/8^* astrocytes, BMAL1 binding is extensively redistributed and specifically decoupled from a coherent set of differentiation-associated targets, while global promoter accessibility changes do not explain which sites are gained or lost. Instead, WT-specific enrichment of motifs corresponding to known circadian co-binder classes points to cooperative recruitment as a key mechanistic lever. More broadly, MACS-CC provides a generalizable approach to map TF occupancy in glial populations *in vivo*, enabling future work to dissect how partner networks, inflammatory signaling, and metabolic state converge on circadian TF targeting in neurodegeneration. Finally, in the context of circadian-based therapeutic interest, our data argue that targeting chromatin accessibility alone may be insufficient to modulate disease-altered clock TF binding; instead, interventions that stabilize or restore cooperative factor interactions may offer a more plausible route to re-establishing functional circadian transcriptional programs in CLN3-deficient astrocytes. It will be important to determine whether these fundamental observations extend to other disorders in which circadian dysregulation and progressive neuron loss occur.

## ACKNOWLEDGEMENTS

We thank the High Throughput Computing Facility @ CGS_SB for computational resources. We thank the Hope Center Viral Vectors Core for AAV production and the Washington University Genome Technology Access Center (GTAC) and Jessica Hoisington-Lopez and Maria-Lynn Crosby at WashU’s DNA Sequencing Innovation Lab (DSIL) for sequencing. GTAC also provided RNA-seq library preparation services. We thank Pat Sheehan (Musiek laboratory) for early discussions that helped initiate the central idea for this study. We also thank members in the Mitra laboratory for providing valuable suggestions.

## AUTHOR CONTRIBUTIONS

India H. Reiss: Conceptualization, Investigation, Formal analysis, Methodology, Validation, Visualization, Writing - original draft. Jonathan D. Cooper: Conceptualization, Resources, Writing - review & editing. Erik S. Musiek: Conceptualization, Writing - review & editing. Robi David Mitra: Conceptualization, Project administration, Funding acquisition, Resources, Supervision, Writing - review & editing.

## SUPPLEMENTARY DATA

Supplementary Data are available at NAR online.

## CONFLICT OF INTEREST

R.D.M. is listed as an inventor on US patent US20200181626A1.

## FUNDING

This work was supported by National Institutes of Health BRAIN Initiative grant [1RF1MH117070], the National Institute of Mental Health [RM1MH138313], the National Institute of Dental and Craniofacial Research [R01DE032865], a National Institutes of Health / National Institute on Aging F31 Predoctoral Fellowship [F31AG076286], and the Goldfarb Center for Neuronal Resiliency.

## DATA AVAILABILITY

Raw sequencing data have been deposited in the NCBI Sequence Read Archive (SRA) under BioProject PRJNA1419685 and in the NIH GEO repository under GEO accession numbers GSE318942 and GSE329982.

## Notes

### Summary of Updates

Changed format to match NAR guidelines (word limit on abstract, Materials and Methods listed before Results section, etc.).

## REFERENCES

1. Khakh, B.S. and Sofroniew, M.V. (2015) Diversity of astrocyte functions and phenotypes in neural circuits. Nat. Neurosci., 18, 942–952.

2. Brandebura, A.N., Paumier, A., Onur, T.S. and Allen, N.J. (2023) Astrocyte contribution to dysfunction, risk and progression in neurodegenerative disorders. Nat. Rev. Neurosci., 24, 23–39.

3. Musiek, E.S., Xiong, D.D. and Holtzman, D.M. (2015) Sleep, circadian rhythms, and the pathogenesis of Alzheimer disease. Exp. Mol. Med., 47, e148.

4. Edison, P. (2024) Astroglial activation: Current concepts and future directions. Alzheimers. Dement., 20, 3034–3053.

5. Sofroniew, M.V. (2014) Astrogliosis. Cold Spring Harb. Perspect. Biol., 7, a020420.

6. Price, B.R., Johnson, L.A. and Norris, C.M. (2021) Reactive astrocytes: The nexus of pathological and clinical hallmarks of Alzheimer’s disease. Ageing Res. Rev., 68, 101335.

7. Lananna, B.V., Nadarajah, C.J., Izumo, M., Cedeño, M.R., Xiong, D.D., Dimitry, J., Tso, C.F., McKee, C.A., Griffin, P., Sheehan, P.W., et al. (2018) Cell-autonomous regulation of astrocyte activation by the circadian clock protein BMAL1. Cell Rep., 25, 1–9.e5.

8. Sheehan, P.W., Fass, S.B., Sapkota, D., Kang, S., Hollis, H.C., Lawrence, J.H., Park, S., Sharma, A., Schafer, D.P., Anafi, R.C., et al. (2025) A glial circadian gene expression atlas reveals cell-type and disease-specific reprogramming in response to amyloid pathology or aging. Nat. Neurosci., 28, 2366–2379.

9. Zhang, R., Lahens, N.F., Ballance, H.I., Hughes, M.E. and Hogenesch, J.B. (2014) A circadian gene expression atlas in mammals: implications for biology and medicine. Proc. Natl. Acad. Sci. U. S. A., 111, 16219–16224.

10. Eckel-Mahan, K.L., Patel, V.R., de Mateo, S., Orozco-Solis, R., Ceglia, N.J., Sahar, S., Dilag-Penilla, S.A., Dyar, K.A., Baldi, P. and Sassone-Corsi, P. (2013) Reprogramming of the circadian clock by nutritional challenge. Cell, 155, 1464–1478.

11. Levine, D.C., Hong, H., Weidemann, B.J., Ramsey, K.M., Affinati, A.H., Schmidt, M.S., Cedernaes, J., Omura, C., Braun, R., Lee, C., et al. (2020) NAD+ controls circadian reprogramming through PER2 nuclear translocation to counter aging. Mol. Cell, 78, 835–849.e7.

12. Montellier, E., Vial, G., Bouyon, S., Ashtiani, K.C., Abdelkarim, S., Lemarie, E., Boutin, A., Kinouchi, K., Baldi, P., Pépin, J.-L., et al. (2026) Chronic intermittent hypoxia reshapes circadian metabolic architecture in a model of sleep apnea. Sci. Adv., 12, eaeb3756.

13. Takahashi, J.S. (2017) Transcriptional architecture of the mammalian circadian clock. Nat. Rev. Genet., 18, 164–179.

14. Yang, G., Chen, L., Grant, G.R., Paschos, G., Song, W.-L., Musiek, E.S., Lee, V., McLoughlin, S.C., Grosser, T., Cotsarelis, G., et al. (2016) Timing of expression of the core clock gene Bmal1 influences its effects on aging and survival. Sci. Transl. Med., 8, 324ra16.

15. Lananna, B.V. and Musiek, E.S. (2020) The wrinkling of time: Aging, inflammation, oxidative stress, and the circadian clock in neurodegeneration. Neurobiol. Dis., 139, 104832.

16. Colwell, C.S. (2021) Defining circadian disruption in neurodegenerative disorders. J. Clin. Invest., 131.

17. Namgyal, D. and Lim, C.-S. (2025) Circadian rhythm dysfunction in neurodegenerative diseases: A bidirectional perspective and therapeutic potential. Nat. Sci. Sleep, 17, 2969–2989.

18. Brent, M.R. (2016) Past roadblocks and new opportunities in transcription factor network mapping. Trends Genet., 32, 736–750.

19. Ziółkowska, E.A., Takahashi, K., Dickson, P.I., Sardiello, M., Sands, M.S. and Cooper, J.D. (2025) Neuronal ceroid lipofuscinosis: underlying mechanisms and emerging therapeutic targets. Nat. Rev. Neurol., 21, 606–622.

20. Takahashi, K., Nelvagal, H.R., Lange, J. and Cooper, J.D. (2022) Glial dysfunction and its contribution to the pathogenesis of the neuronal ceroid lipofuscinoses. Front. Neurol., 13, 886567.

21. Parviainen, L., Dihanich, S., Anderson, G.W., Wong, A.M., Brooks, H.R., Abeti, R., Rezaie, P., Lalli, G., Pope, S., Heales, S.J., et al. (2017) Glial cells are functionally impaired in juvenile neuronal ceroid lipofuscinosis and detrimental to neurons. Acta Neuropathol. Commun., 5, 74.

22. Johnson, T.B., Brudvig, J.J., Likhite, S., Pratt, M.A., White, K.A., Cain, J.T., Booth, C.D., Timm, D.J., Davis, S.S., Meyerink, B., et al. (2023) Early postnatal administration of an AAV9 gene therapy is safe and efficacious in CLN3 disease. Front. Genet., 14, 1118649.

23. Pontikis, C.C., Cotman, S.L., MacDonald, M.E. and Cooper, J.D. (2005) Thalamocortical neuron loss and localized astrocytosis in the Cln3Deltaex7/8 knock-in mouse model of Batten disease. Neurobiol. Dis., 20, 823–836.

24. Cotman, S.L., Vrbanac, V., Lebel, L.-A., Lee, R.L., Johnson, K.A., Donahue, L.-R., Teed, A.M., Antonellis, K., Bronson, R.T., Lerner, T.J., et al. (2002) Cln3(Deltaex7/8) knock-in mice with the common JNCL mutation exhibit progressive neurologic disease that begins before birth. Hum. Mol. Genet., 11, 2709–2721.

25. Lerner, T.J., Boustany, R.-M.N., Anderson, J.W., D’Arigo, K.L., Schlumpf, K., Buckler, A.J., Gusella, J.F. and Haines, J.L. (1995) Isolation of a novel gene underlying Batten disease, CLN3. The International Batten Disease Consortium. Cell, 82, 949–957.

26. Yen, A., Mateusiak, C., Sarafinovska, S., Gachechiladze, M.A., Guo, J., Chen, X., Moudgil, A., Cammack, A.J., Hoisington-Lopez, J., Crosby, M., et al. (2023) Calling Cards: A customizable platform to longitudinally record protein-DNA interactions over time in cells and tissues. Curr. Protoc., 3, e883.

27. Yamazaki, A., Shue, F., Yamazaki, Y., Martens, Y.A., Bu, G. and Liu, C.-C. (2021) Preparation of single cell suspensions enriched in mouse brain vascular cells for single-cell RNA sequencing. STAR Protoc., 2, 100715.

28. Miltenyi, S., Müller, W., Weichel, W. and Radbruch, A. (1990) High gradient magnetic cell separation with MACS. Cytometry, 11, 231–238.

29. Lalli, M., Yen, A., Thopte, U., Dong, F., Moudgil, A., Chen, X., Milbrandt, J., Dougherty, J.D. and Mitra, R.D. (2022) Measuring transcription factor binding and gene expression using barcoded self-reporting transposon calling cards and transcriptomes. NAR Genom. Bioinform., 4, lqac061.

30. Moudgil, A., Wilkinson, M.N., Chen, X., He, J., Cammack, A.J., Vasek, M.J., Lagunas, T., Jr, Qi, Z., Lalli, M.A., Guo, C., et al. (2020) Self-reporting transposons enable simultaneous readout of gene expression and transcription factor binding in single cells. Cell, 182, 992–1008.e21.

31. Cammack, A.J., Moudgil, A., Chen, J., Vasek, M.J., Shabsovich, M., McCullough, K., Yen, A., Lagunas, T., Maloney, S.E., He, J., et al. (2020) A viral toolkit for recording transcription factor-DNA interactions in live mouse tissues. Proc. Natl. Acad. Sci. U. S. A., 117, 10003–10014.

32. Wang, H., Mayhew, D., Chen, X., Johnston, M. and Mitra, R.D. (2012) ‘Calling cards’ for DNA-binding proteins in mammalian cells. Genetics, 190, 941–949.

33. Marsh, S.E., Walker, A.J., Kamath, T., Dissing-Olesen, L., Hammond, T.R., de Soysa, T.Y., Young, A.M.H., Murphy, S., Abdulraouf, A., Nadaf, N., et al. (2022) Dissection of artifactual and confounding glial signatures by single-cell sequencing of mouse and human brain. Nat. Neurosci., 25, 306–316.

34. van den Brink, S.C., Sage, F., Vértesy, Á., Spanjaard, B., Peterson-Maduro, J., Baron, C.S., Robin, C. and van Oudenaarden, A. (2017) Single-cell sequencing reveals dissociation-induced gene expression in tissue subpopulations. Nat. Methods, 14, 935–936.

35. Hochbaum, D.R., Hulshof, L., Urke, A., Wang, W., Dubinsky, A.C., Farnsworth, H.C., Hakim, R., Lin, S., Kleinberg, G., Robertson, K., et al. (2024) Thyroid hormone remodels cortex to coordinate body-wide metabolism and exploration. Cell, 187, 5679–5697.e23.

36. Burkovetskaya, M., Karpuk, N., Xiong, J., Bosch, M., Boska, M.D., Takeuchi, H., Suzumura, A. and Kielian, T. (2014) Evidence for aberrant astrocyte hemichannel activity in Juvenile Neuronal Ceroid Lipofuscinosis (JNCL). PLoS One, 9, e95023.

37. Ali, A.A.H., Schwarz-Herzke, B., Mir, S., Sahlender, B., Victor, M., Görg, B., Schmuck, M., Dach, K., Fritsche, E., Kremer, A., et al. (2019) Deficiency of the clock gene Bmal1 affects neural progenitor cell migration. Brain Struct. Funct., 224, 373–386.

38. Malik, A., Kondratov, R.V., Jamasbi, R.J. and Geusz, M.E. (2015) Circadian clock genes are essential for normal adult neurogenesis, differentiation, and fate determination. PLoS One, 10, e0139655.

39. Bering, T., Gadgaard, C., Vorum, H., Honoré, B. and Rath, M.F. (2023) Diurnal proteome profile of the mouse cerebral cortex: Conditional deletion of the Bmal1 circadian clock gene elevates astrocyte protein levels and cell abundance in the neocortex and hippocampus. Glia, 71, 2623–2641.

40. Beytebiere, J.R., Trott, A.J., Greenwell, B.J., Osborne, C.A., Vitet, H., Spence, J., Yoo, S.-H., Chen, Z., Takahashi, J.S., Ghaffari, N., et al. (2019) Tissue-specific BMAL1 cistromes reveal that rhythmic transcription is associated with rhythmic enhancer-enhancer interactions. Genes Dev., 33, 294–309.

41. Liu, C., Wang, M., Wei, X., Wu, L., Xu, J., Dai, X., Xia, J., Cheng, M., Yuan, Y., Zhang, P., et al. (2019) An ATAC-seq atlas of chromatin accessibility in mouse tissues. Sci. Data, 6, 65.

42. Hong, H.-K., Maury, E., Ramsey, K.M., Perelis, M., Marcheva, B., Omura, C., Kobayashi, Y., Guttridge, D.C., Barish, G.D. and Bass, J. (2018) Requirement for NF-κB in maintenance of molecular and behavioral circadian rhythms in mice. Genes Dev., 32, 1367–1379.

43. Leung, A., Trac, C., Du, J., Natarajan, R. and Schones, D.E. (2016) Persistent chromatin modifications induced by high fat diet. J. Biol. Chem., 291, 10446–10455.

44. Levine, D.C., Kuo, H.-Y., Hong, H.-K., Cedernaes, J., Hepler, C., Wright, A.G., Sommars, M.A., Kobayashi, Y., Marcheva, B., Gao, P., et al. (2021) NADH inhibition of SIRT1 links energy state to transcription during time-restricted feeding. Nat. Metab., 3, 1621–1632.

45. Fan, Y., Qian, H., Zhang, M., Tao, C., Li, Z., Yan, W., Huang, Y., Zhang, Y., Xu, Q., Wang, X., et al. (2023) Caloric restriction remodels the hepatic chromatin landscape and bile acid metabolism by modulating the gut microbiota. Genome Biol., 24, 98.

46. Lange, J., Haslett, L.J., Lloyd-Evans, E., Pocock, J.M., Sands, M.S., Williams, B.P. and Cooper, J.D. (2018) Compromised astrocyte function and survival negatively impact neurons in infantile neuronal ceroid lipofuscinosis. Acta Neuropathol. Commun., 6, 74.

47. Bosch, M.E. and Kielian, T. (2019) Astrocytes in juvenile neuronal ceroid lipofuscinosis (CLN3) display metabolic and calcium signaling abnormalities. J. Neurochem., 148, 612–624.

48. Yasa, S., Butz, E.S., Colombo, A., Chandrachud, U., Montore, L., Tschirner, S., Prestel, M., Sheridan, S.D., Müller, S.A., Groh, J., et al. (2024) Loss of CLN3 in microglia leads to impaired lipid metabolism and myelin turnover. *Commun*. Biol., 7, 1373.

49. Xiong, J. and Kielian, T. (2013) Microglia in juvenile neuronal ceroid lipofuscinosis are primed toward a pro-inflammatory phenotype. J. Neurochem., 127, 245–258.

50. Berve, K., West, B.L., Martini, R. and Groh, J. (2020) Sex- and region-biased depletion of microglia/macrophages attenuates CLN1 disease in mice. J. Neuroinflammation, 17, 323.

51. Cooper, J.D., Tarczyluk, M.A. and Nelvagal, H.R. (2015) Towards a new understanding of NCL pathogenesis. Biochim. Biophys. Acta, 1852, 2256–2261.

52. Nelvagal, H.R., Lange, J., Takahashi, K., Tarczyluk-Wells, M.A. and Cooper, J.D. (2020) Pathomechanisms in the neuronal ceroid lipofuscinoses. Biochim. Biophys. Acta Mol. Basis Dis., 1866, 165570.

53. Pontikis, C.C., Cella, C.V., Parihar, N., Lim, M.J., Chakrabarti, S., Mitchison, H.M., Mobley, W.C., Rezaie, P., Pearce, D.A. and Cooper, J.D. (2004) Late onset neurodegeneration in the Cln3-/- mouse model of juvenile neuronal ceroid lipofuscinosis is preceded by low level glial activation. Brain Res., 1023, 231–242.

54. Kielar, C., Maddox, L., Bible, E., Pontikis, C.C., Macauley, S.L., Griffey, M.A., Wong, M., Sands, M.S. and Cooper, J.D. (2007) Successive neuron loss in the thalamus and cortex in a mouse model of infantile neuronal ceroid lipofuscinosis. Neurobiol. Dis., 25, 150–162.

55. Partanen, S., Haapanen, A., Kielar, C., Pontikis, C., Alexander, N., Inkinen, T., Saftig, P., Gillingwater, T.H., Cooper, J.D. and Tyynelä, J. (2008) Synaptic changes in the thalamocortical system of cathepsin D-deficient mice: a model of human congenital neuronal ceroid-lipofuscinosis: A model of human congenital neuronal ceroid-lipofuscinosis. J. Neuropathol. Exp. Neurol., 67, 16–29.

56. Zheng, Y., Pan, L., Wang, F., Yan, J., Wang, T., Xia, Y., Yao, L., Deng, K., Zheng, Y., Xia, X., et al. (2023) Neural function of Bmal1: an overview. Cell Biosci., 13, 1.

57. Bouteldja, A.A., Penichet, D., Srivastava, L.K. and Cermakian, N. (2024) The circadian system: A neglected player in neurodevelopmental disorders. Eur. J. Neurosci., 60, 3858–3890.

58. Ali, A.A.H., Schwarz-Herzke, B., Rollenhagen, A., Anstötz, M., Holub, M., Lübke, J., Rose, C.R., Schnittler, H.-J. and von Gall, C. (2020) Bmal1-deficiency affects glial synaptic coverage of the hippocampal mossy fiber synapse and the actin cytoskeleton in astrocytes. Glia, 68, 947–962.

59. Gomez-Giro, G., Arias-Fuenzalida, J., Jarazo, J., Zeuschner, D., Ali, M., Possemis, N., Bolognin, S., Halder, R., Jäger, C., Kuper, W.F.E., et al. (2019) Synapse alterations precede neuronal damage and storage pathology in a human cerebral organoid model of CLN3-juvenile neuronal ceroid lipofuscinosis. Acta Neuropathol. Commun., 7, 222.

60. Singh, J.B., Burris, D.M., Bhuyan, S., Thurston, T., Oschmann, A., Jankowski, C., Lu, W., Rabinowitz, J.D. and Ahrens-Nicklas, R.C. (2025) CLN3 disease disrupts very early postnatal hippocampal maturation. Sci. Rep., 15, 24411.

61. Barnat, M., Capizzi, M., Aparicio, E., Boluda, S., Wennagel, D., Kacher, R., Kassem, R., Lenoir, S., Agasse, F., Braz, B.Y., et al. (2020) Huntington’s disease alters human neurodevelopment. Science, 369, 787–793.

62. Finger, E., Malik, R., Bocchetta, M., Coleman, K., Graff, C., Borroni, B., Masellis, M., Laforce, R., Greaves, C.V., Russell, L.L., et al. (2023) Neurodevelopmental effects of genetic frontotemporal dementia in young adult mutation carriers. Brain, 146, 2120–2131.

63. Hendricks, E., Quihuis, A.M., Hung, S.-T., Chang, J., Dorjsuren, N., Der, B., Staats, K.A., Shi, Y., Sta Maria, N.S., Jacobs, R.E., et al. (2023) The C9ORF72 repeat expansion alters neurodevelopment. Cell Rep., 42, 112983.

64. Wulansari, N., Darsono, W.H.W., Woo, H.-J., Chang, M.-Y., Kim, J., Bae, E.-J., Sun, W., Lee, J.-H., Cho, I.-J., Shin, H., et al. (2021) Neurodevelopmental defects and neurodegenerative phenotypes in human brain organoids carrying Parkinson’s disease-linked DNAJC6 mutations. Sci. Adv., 7, eabb1540.

65. Schwamborn, J.C. (2018) Is Parkinson’s disease a neurodevelopmental disorder and will brain organoids help us to understand it? Stem Cells Dev., 27, 968–975.

66. Meyer-Acosta, K.K., Diaz-Guerra, E., Varma, P., Aruk, A., Mirsadeghi, S., Muniz-Perez, A., Rafati, Y., Hosseini, A., Nieto-Estevez, V., Giugliano, M., et al. (2025) APOE4 impacts cortical neurodevelopment and alters network formation in human brain organoids. Stem Cell Reports, 20, 102537.

67. Holmes, A.G., Parker, J.B., Sagar, V., Truica, M.I., Soni, P.N., Han, H., Schiltz, G.E., Abdulkadir, S.A. and Chakravarti, D. (2022) A MYC inhibitor selectively alters the MYC and MAX cistromes and modulates the epigenomic landscape to regulate target gene expression. Sci. Adv., 8, eabh3635.

68. Hong, S. and Song, M.-R. (2014) STAT3 but not STAT1 is required for astrocyte differentiation. PLoS One, 9, e86851.

69. Ceyzériat, K., Abjean, L., Carrillo-de Sauvage, M.-A., Ben Haim, L. and Escartin, C. (2016) The complex STATes of astrocyte reactivity: How are they controlled by the JAK-STAT3 pathway? Neuroscience, 330, 205–218.

70. Nakashima, K., Yanagisawa, M., Arakawa, H., Kimura, N., Hisatsune, T., Kawabata, M., Miyazono, K. and Taga, T. (1999) Synergistic signaling in fetal brain by STAT3-Smad1 complex bridged by p300. Science, 284, 479–482.

71. Itoh, Y., Saitoh, M. and Miyazawa, K. (2018) Smad3-STAT3 crosstalk in pathophysiological contexts. Acta Biochim. Biophys. Sin. (Shanghai), 50, 82–90.

72. Solt, L.A., Kojetin, D.J. and Burris, T.P. (2011) The REV-ERBs and RORs: molecular links between circadian rhythms and lipid homeostasis. Future Med. Chem., 3, 623–638.

73. Bugge, A., Feng, D., Everett, L.J., Briggs, E.R., Mullican, S.E., Wang, F., Jager, J. and Lazar, M.A. (2012) Rev-erbα and Rev-erbβ coordinately protect the circadian clock and normal metabolic function. Genes Dev., 26, 657–667.

74. Qu, M., Qu, H., Jia, Z. and Kay, S.A. (2021) HNF4A defines tissue-specific circadian rhythms by beaconing BMAL1::CLOCK chromatin binding and shaping the rhythmic chromatin landscape. Nat. Commun., 12, 6350.

75. Bonjoch, J., Solá, P., Reina, O., Mortimer, T., Miroshnikova, Y.A., Wickström, S.A., Stephan-Otto Attolini, C., Álvarez, L., Aznar Benitah, S. and Solanas, G. (2025) BMAL1 and YAP cooperate to hijack enhancers and promote inflammation in the aged epidermis. bioRxiv, 10.1101/2025.04.22.649967.

