## Supplemental Tables for "*In vivo* BMAL1 occupancy mapping using MACS-Calling Cards reveals disease-associated retargeting in *Cln3^Δex7/8^* astrocytes"

Supplementary Tables:

**Supplementary Table S1**: *In vitro* Calling Card plasmids:

| Mitra Lab Plasmid Number | Plasmid Name | Description |
|---|---|---|
| pRM2097 | cmv_mbmal1_hyPBase | Mouse BMAL1-hyPBase transposase fusion for cell line calling card experiments. |
| pRM1114 | pCMV_hyPBase_Centrip | Unfused hyPBase transposase fusion for cell line calling card experiments. |
| pRM1892 | Barcoded PB_SRT_Puro | Barcoded piggyBac transposon vectors for cell line calling card experiments. Reporter is pac (puromycin resistance). All barcodes were equally pooled. |

**Supplementary Table S2**: Genotyping plasmids:

| Genotyping Primer Name | Genotyping Target | Primer Sequence |
|---|---|---|
| B-actin_F | Actb | AGAGGGAAATCGTGCGTGAC |
| B-actin_R | Actb | CAATAGTGATGACCTGGCCGT |
| Egfp_for | Egfp | CCTACGGCGTGCAGTGCTTCAGC |
| Egfp_rev | Egfp | CGGCGAGCTGCACGCTGCGTCCTC |

**Supplementary Table S3**: *In vivo* Calling Card plasmids.

| Mitra Lab Plasmid Number | Plasmid Name | Description |
|---|---|---|
| pRM2095 | AAV_cag_mBMAL1_hyPBase | Mouse BMAL1-hyPBase transposase fusion for AAV calling card experiments. |
| pRM2096 | AAV_cag_hyPBase | Unfused hyPBase transposase fusion for AAV |

| | | |
|---|---|---|
| | | calling card experiments. |
| pRM1891 | Barcoded AAVtdTomato_SRT | Barcoded piggyBac transposon vectors for AAV calling card experiments. The reporter is tdtomato. All barcodes were equally pooled. |

**Supplementary Table S4:** MNase-seq primer and adapter sequences:

| Name | Type | Sequence (example index=underlined) |
|---|---|---|
| MNase1_[index] | Primer | CAAGCAGAAGACGGCATACGAGATCTGTGTAACGGTGACTGGAGTTCAGACGTGTGCTCTTCCGA |
| MNase2_[index] | Primer | AATGATACGGCGACCACCGAGATCTACACACAACGACAAACACTCTTTCCCTACACGACGCTCTTCCGATCT |
| A1 Adapter | Adapter | ACACTCTTTCCCTACACGACGCTCTTCCGATCT |
| A2 Adapter | Adapter | P-GATCGGAAGAGCACACGTCTGAACTCCAGTCAC |

**Supplementary Table S5:** Calling Cards primer sequences:

| Primer Name | Sequence (example index=underlined) |
|---|---|
| SMART_dT18VN | AAGCAGTGGTATCAACGCAGAGTACGTTTTTTTTTTTTTTTTTTTTTTTTTTTTTTVN |
| SRT_PAC_F1 | CAACCTCCCCTTCTACGAGC |
| SRT_tdTomato_F1 | TCCTGTACGGCATGGACGAG |
| SMART | AAGCAGTGGTATCAACGCAGAGT |
| OM_PB_Mitra[index]_mutex_TCG | AATGATACGGCGACCACCGAGATCTACACACAACGACAAACACTCTTTCCCTACACGACGCTCTTCCGATCTTCGTgcgtcaattttacgcagactatcttt |
| Nextera_[index]-10bp | CAAGCAGAAGACGGCATACGAGATGCACT |

| | |
|---|---|
| | GTGCTGTCTCGTGGGCTCGG |

**Supplementary Table S6:** Genes included in Figure 7D:

| Ensembl ID | Gene Name |
|---|---|
| ENSMUSG00000002147 | Stat6 |
| ENSMUSG00000004043 | Stat5a |
| ENSMUSG00000020919 | Stat5b |
| ENSMUSG00000004040 | Stat3 |
| ENSMUSG00000026104 | Stat1 |
| ENSMUSG00000040033 | Stat2 |
| ENSMUSG00000062939 | Stat4 |
| ENSMUSG00000055320 | Tead1 |
| ENSMUSG00000030796 | Tead2 |
| ENSMUSG00000002249 | Tead3 |
| ENSMUSG00000030353 | Tead4 |
| ENSMUSG00000003949 | Hlf |
| ENSMUSG00000056749 | Nfil3 |
| ENSMUSG00000021381 | Barx1 |
| ENSMUSG00000032033 | Barx2 |
| ENSMUSG00000032035 | Ets1 |
| ENSMUSG00000036461 | Elf1 |
| ENSMUSG00000009406 | Elk1 |
| ENSMUSG00000008976 | Gabpa |
| ENSMUSG00000016087 | Fli1 |
| ENSMUSG00000040732 | Erg |
| ENSMUSG00000042414 | Prdm14 |
| ENSMUSG00000014039 | Prdm15 |
| ENSMUSG00000037992 | Rara |
| ENSMUSG00000017491 | Rarb |
| ENSMUSG00000001288 | Rarg |
| ENSMUSG00000015846 | Rxra |
| ENSMUSG00000039656 | Rxrb |
| ENSMUSG00000015843 | Rxrg |
| ENSMUSG00000022383 | Ppara |
| ENSMUSG00000002250 | Ppard |
| ENSMUSG00000000440 | Pparg |
| ENSMUSG00000058756 | Thra |
| ENSMUSG00000021779 | Thrb |
| ENSMUSG00000022479 | Vdr |
| ENSMUSG00000060601 | Nr1h2 |

| | |
|---|---|
| ENSMUSG00000002108 | Nr1h3 |
| ENSMUSG00000047638 | Nr1h4 |
| ENSMUSG00000022809 | Nr1i2 |
| ENSMUSG00000005677 | Nr1i3 |
| ENSMUSG00000017950 | Hnf4a |
| ENSMUSG00000017688 | Hnf4g |
| ENSMUSG00000069171 | Nr2f1 |
| ENSMUSG00000030551 | Nr2f2 |
| ENSMUSG00000002393 | Nr2f6 |
| ENSMUSG00000020889 | Nr1d1 |
| ENSMUSG00000021775 | Nr1d2 |
| ENSMUSG00000032238 | Rora |
| ENSMUSG00000036192 | Rorb |
| ENSMUSG00000028150 | Rorc |
| ENSMUSG00000024955 | Esrra |
| ENSMUSG00000021255 | Esrrb |
| ENSMUSG00000026610 | Esrrg |
| ENSMUSG00000030067 | Foxp1 |
| ENSMUSG00000029563 | Foxp2 |
| ENSMUSG00000039521 | Foxp3 |
| ENSMUSG00000023991 | Foxp4 |
| ENSMUSG00000035451 | Foxa1 |
| ENSMUSG00000037025 | Foxa2 |
| ENSMUSG00000040891 | Foxa3 |
| ENSMUSG00000044167 | Foxo1 |
| ENSMUSG00000048756 | Foxo3 |
| ENSMUSG00000042903 | Foxo4 |
