## Supplemental Figures for "*In vivo* BMAL1 occupancy mapping using MACS-Calling Cards reveals disease-associated retargeting in *Cln3^Δex7/8^* astrocytes"

Supplementary Figures:

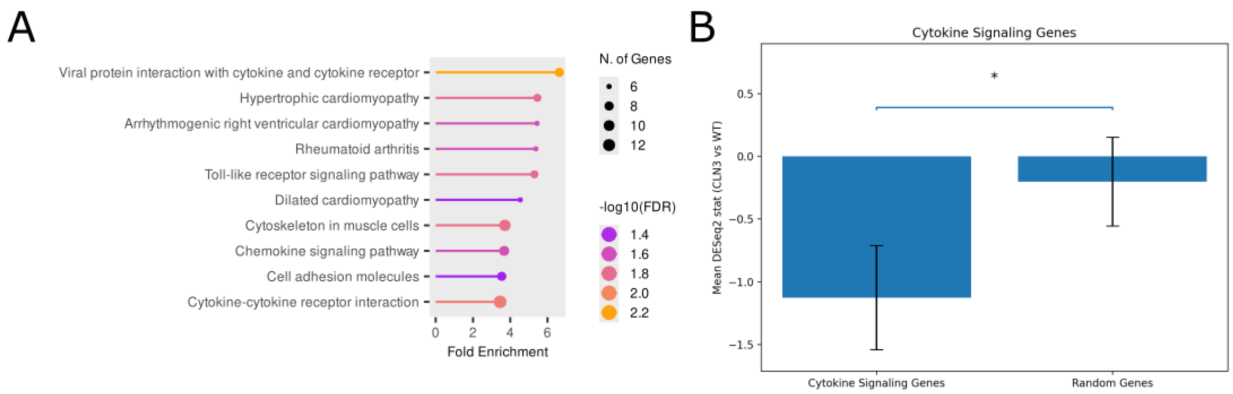


**Supplementary Figure S1**: Differentially expressed genes are enriched for cytokine signaling pathways. (A) Pathway enrichment analysis of DEGs performed in ShinyGO identifies cytokine-related pathways among the most significantly enriched terms (e.g., cytokine-cytokine receptor interaction, chemokine signaling, Toll-like receptor signaling). Dot position indicates fold enrichment; dot size indicates the number of DEGs in each pathway; color denotes significance (-log10 FDR). (B) Cytokine signaling genes show a larger average DESeq2 stat value (CLN3 vs WT) than a randomly selected gene set ($P = 0.005$, two sided permutation test).

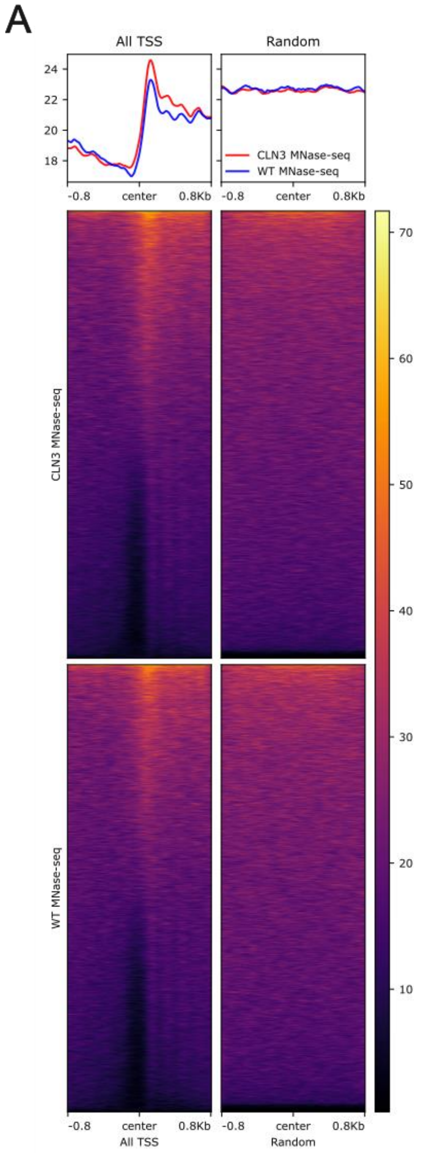


**Supplementary Figure S2**: MNase-seq quality control in WT and *Cln3*$^{\Delta ex7/8}$ astrocytes. (A) Aggregate MNase-seq profiles (top) and corresponding heatmaps (bottom) centered on annotated transcription start sites (All TSS; ±0.8 kb) reveal a pronounced nucleosome-free region (NFR) at the TSS (center) flanked by phased nucleosomes in both WT (blue) and *Cln3*$^{\Delta ex7/8}$ (red) datasets. In contrast, profiles centered on random genomic positions show no organized depletion or phasing. Heatmaps display MNase-seq signal around each site for *Cln3*$^{\Delta ex7/8}$ (upper) and WT (lower); color indicates MNase-seq signal intensity (scale at right).
